# Synergistic Inhibition of Notch Signaling and Forced Cell Cycle Re-entry Drive Müller Glia Reprogramming in Uninjured Mouse Retina

**DOI:** 10.64898/2026.03.09.710684

**Authors:** Baoshan Liao, Chengshang Lyu, Yuqing Jiang, Shanggong Liu, Waiho Wong, Jiadong Zhang, Hoyin Tsang, Junxi Xie, Lingxi Chen, Qinrong Zhang, Wenjun Xiong

**Author notes:** Corresponding author: Wenjun Xiong, Address: The Department of Biomedical Sciences, City University of Hong Kong, Kowloon, Hong Kong, China.

## Abstract

In regenerative species, such as teleost fish, Müller glia (MG) autonomously re-enter the cell cycle after injury and give rise to functional retinal neurons. In contrast, the loss of retinal neurons in mammals is irreversible due to the limited proliferative and regenerative ability of MG. Various strategies have been developed to induce proliferation of mature mouse MG with or without injury, yet most MG daughter cells retain glial cell fate. Here, we found that MG progenies maintain high Notch signaling, which may constrain their neurogenic potential. Conditional deletion of *Rbpj*, the central transcriptional effector of Notch, induced limited MG-to-neuron conversion in mature MG without proliferation. However, *Rbpj* deletion, combined with forced MG proliferation by overexpressing *cyclin D1* and suppressing *p27^Kip1^*, significantly promoted MG dedifferentiation and ectopic expression of the neuronal marker Otx2 in MG daughter cells in uninjured mouse retina. Combining Notch inhibition with MG cell cycle re-activation not only increased the numbers of bipolar- and amacrine-like cells generated from MG but also promoted the further differentiation toward ON-cone, OFF-cone, and rod-bipolar subtypes. Single-nucleus RNA and ATAC sequencing data revealed that Notch inhibition facilitated the formation of MG-derived progenitor-like cells while MG proliferation increased chromatin accessibility of neurogenic genes. Notably, most MG-derived cells survived long term despite incomplete maturation. Together, our findings delineate how Notch inhibition and MG proliferation, alone or in combination, influence the regenerative potential of MG in the mammalian retina.

## Introduction

Müller glia share a common lineage with retinal neurons, serving as the only glial cell type differentiated from retinal progenitor cells (RPCs) while retaining a transcriptomic profile similar to their progenitors(*1–3*). Beyond their role in maintaining retinal homeostasis, MG function as a latent stem cell population in lower vertebrates. In zebrafish, retinal injury triggers a robust regenerative response wherein quiescent MG dedifferentiate, re-enter the cell cycle, and undergo asymmetric division to self-renew and generate multipotent retinal progenitors. These progenitor cells subsequently differentiate to regenerate all major retinal neuron types, replacing damaged cells(*4–6*). However, this regenerative capacity is progressively restricted across vertebrate evolution. While MG in young chicks can initiate a single mitotic cycle and transition to MG-derived progenitor cells (MGPCs), MGPCs only differentiate into amacrine cells and bipolar cells(*7*, *8*). In mice and humans, MG lack the intrinsic capacity to autonomously re-enter the cell cycle or regenerate lost neurons, representing a major barrier to treating retinal degenerative diseases(*2*, *6*, *9*).

Overcoming MG quiescence is the first step toward regeneration. We previously demonstrated that the high level of *p27^Kip1^*, a cell cycle inhibitor, and the low level of *cyclin D1* in mature mouse MG prevent them from re-entering the cell cycle. Concurrent downregulation of *p27^Kip1^* and upregulation of *cyclin D1*, which was delivered via a single AAV vector termed the Cell Cycle Activator (CCA), synergistically promote MG proliferation. CCA-driven MG proliferation further promoted the dedifferentiation of MG following mitosis, but the vast majority of MG eventually reverted to their glial identity(*10*). Similarly, direct activation of the Wnt/β-catenin signaling pathway or bypassing the Hippo pathway via overexpressing Hippo non-responsive form of YAP (YAP5SA) also led to spontaneous re-entry into the cell cycle and transiently reprogramming into a progenitor cell-like state, but no regeneration of mature neurons was reported(*11–13*). These findings indicate that while cell proliferation is necessary to expand the MG pool and initiate dedifferentiation, it is insufficient to drive functional neurogenesis, implying that the existence of additional molecular barriers that enforce glial identity.

Notch signaling represents a primary candidate for this barrier. During development, Notch acts as a binary switch, maintaining the RPC pool and specifying glial fate. Downregulation of Notch is required for neuronal differentiation, whereas sustained activity promotes MG formation(*12*, *14*, *15*). After development, this pathway remains active in mature MG, enforcing quiescence in the uninjured retina(*14*, *16*). In zebrafish, injury induces the downregulation of Notch receptors and downstream target genes *Hes*/*Hey* family, thereby derepressing the proneural genes *Ascl1* to promote neurogenesis(*6*, *14*, *17–19*). Conversely, in mammalian central nervous system, Notch signaling actively suppresses neurogenesis in contexts ranging from the cortex to the cochlea(*20–23*). Recent studies in the mouse retina have shown that disrupting Notch, particularly in combination with knockout of *Nuclear factor I a/b/x* (*Nfi a/b/x*) or overexpressing the Yamanaka factor, *octamer-binding transcription factor 4* (*Oct4*), enhance MG reprogramming efficiency(*24*, *25*).

In this study, we investigated why postmitotic MG fail to yield neurons. We found that Notch signaling remains active in the postmitotic MG-derived cells, which may prevent their diversion toward a neuronal fate. We hypothesized that inhibition of Notch signaling would unlock the neurogenic potential of proliferating MG. By combining CCA treatment with MG-specific deletion of the Notch transcriptional effector, *Recombination signal binding protein for immunoglobulin kappa J region* (*Rbpj*), we achieved robust reprogramming of MG into bipolar- and amacrine-like neurons in uninjured adult mouse retina. Single-nucleus RNA sequencing (snRNA-seq) and RNA *in situ* hybridization revealed that these newborn neurons exhibit a retinal neuron transcriptional signature, expressing markers characteristic of amacrine cells (AC) and subtypes of bipolar cells (BC). Single nucleus ATAC-sequencing (snATAC-seq) analysis demonstrated that CCA treatment induced chromatin opening at key neurogenic genes, a priming event that is functionally capitalized upon by Notch inhibition. Notably, the reprogrammed MG displayed long-term survival up to 9 months post-treatment. Together, our study demonstrated the synergistic effects of cell proliferation and Notch inhibition on MG reprogramming in the absence of retinal injury, providing molecular and temporal insights into this process.

## Results

### CCA-induced MG progeny maintains high Notch signaling

Previously, we developed an AAV_7m8_-GFAP-*cyclinD1*-*p27^Kip1^*shRNA vector, designated as CCA, which enables simultaneous *cyclin D1* overexpression and *p27^Kip1^* knockdown specifically in MG (Fig. 1a) (*10*). When injected intravitreally, this vector achieves near-complete MG transduction near the injection site and drives more than half of the MG population to enter the cell cycle once (*10*). Following mitosis, these MG transiently and partially dedifferentiated, but the vast majority of postmitotic MG revert to their original glial identify by four months post-treatment (*10*). Fewer than 1% MG completely lost MG identity, as shown by negative Sox9 staining and decreased GFP signal (Fig. S1a-b), while expressing high levels of Otx2, a pro-neuronal marker (Fig. S1c).

**Figure 1.**
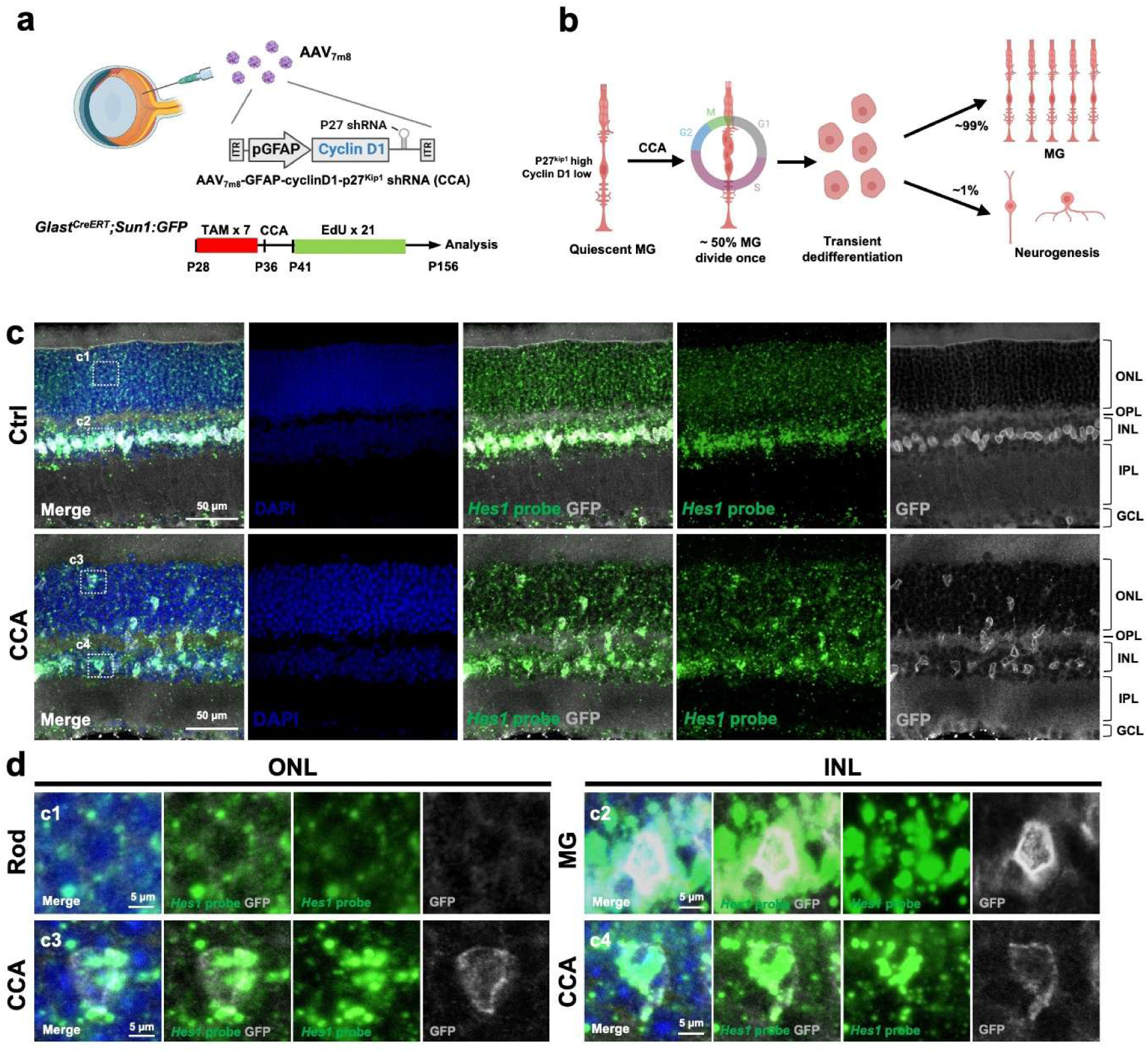
MG and MG-derived cells maintained high level of Notch signaling. (a) Schematic representations of the intravitreal injection of AAV_7m8_-GFAP-*cyclinD1*-*p27^Kip1^* shRNA-WPRE (CCA). (b) The overview process of MG proliferation induced by CCA. (c) *Hes1* mRNA *in situ* hybridization in the control and the *Glast-Cre^ERT^;Sun1:GFP* mouse retinas harvested at four months post CCA injection. (d) Magnified views of the highlighted regions in (c). n=3 mice.

To understand this blockade in neurogenesis, we analyzed the single-cell RNA sequencing (scRNA-seq) data from our previous study(*10*) to characterize the activity of Notch signaling in postmitotic MG. We found that key Notch signaling components, including the receptor *Notch1*, the central regulator *Rbpj*, and the downstream effector *Hes1*, remained highly expressed in postmitotic MG following cell cycle re-activation (Fig. S1d-g). In contrast, these genes were expressed at negligible levels in native retinal neurons, such as rod photoreceptors (Fig S1f-g).

We confirmed this finding using RNA *in situ* hybridization to assess *Hes1* mRNA levels in the *Glast-Cre^ERT^; Sun1:GFP* mice, in which MG nuclei were specifically labelled by nuclear membrane-localized Sun1-tagged GFP. In control retinas, *Hes1* mRNA was abundant in MG within the inner nuclear layer (INL) but minimal in the photoreceptors residing in the outer nuclear layer (ONL) (Fig. 1c-d). In CCA-treated retinas, even in the MG-derived cells that had migrated to the ONL, high *Hes1* mRNA levels maintained (Fig. 1c-d). Given the established role of Notch as a suppressor of neurogenesis, we hypothesized that this persistent Notch activity constitutes the primary barrier preventing CCA-treated MG from differentiating into neurons. Consequently, we reasoned that inhibiting Notch signaling in MG would remove this brake and promote their successful reprogramming into neurons.

### *Rbpj* deletion in late RPC promotes rod genesis at the expense of MG and bipolar cells

*Rbpj* is the essential nuclear mediator of Notch receptors (*26–28*). *Rbpj* knockout in the retina led to efficient inhibition of Notch signaling, as validated by lowered *Hes1* mRNA expression in *Rbpj-/-* cells (Fig. S2a-e). *Glast* is expressed in RPCs in neonatal mice and later restricted to the specified MG(*29*, *30*). Prior to assessing MG-specific *Rbpj* knockout, we first induced *Rbpj* deletion in the late RPCs by administrating Tamoxifen (TAM) to *Glast-Cre^ERT^;Rbpj^flox/flox^;tdTomato(tdT)* mice at postnatal day 1 (P1), then analysed retinas at P12 (Fig. S3a-b). *Rbpj*-/- retinas had a significantly higher proportion of late-born rod photoreceptors, labelled by tdT in the ONL, compared to the control and *Rbpj*+/- retinas, whereas the early-born cone photoreceptors, identified by the mCAR marker, were unaffected (Fig. S3c-d, Fig. S4a-c). Conversely, *Rbpj* deletion significantly impaired the generation of MG (Sox9+) and bipolar cells (Otx2+ in the INL) from late RPCs (Fig. S3c, e, Fig. S5a-c). The generation of amacrine cells, labelled by HuC/D and Pax6 in the INL, and retinal ganglion cells (RGCs), labelled by Rbpms in RGC layer, remained unaffected (Fig. S6a-c, Fig. S7a-c). These developmental data confirm that *Rbpj* removal biases progenitors toward a photoreceptor at the expense of MG and bipolar cells, consistent with the previous report using *Notch1* knockout in the late PRCs(*31*).

### *Rbpj* loss in mature MG triggers inefficient direct glia-to-neuron conversion

We next tested whether Notch inhibition alone is sufficient to reprogram mature MG. We induced *Rbpj* deletion in fully developed retinas at P28 and analysed them at 3 weeks and 4 months post-treatment (Fig. 2a). At 3 weeks, all *Rbpj* KO MG expressed the glial marker Sox9 (Fig. 2b–d). However, by 4 months, approximately 7.6% of GFP-positive MG-derived cells had lost Sox9 and 7.5% had begun expressing the neuronal marker Otx2 (Fig. 2e, Fig. S8a-c). These findings indicate that Notch inhibition alone elicits a slow, progressive dedifferentiation in a subset of MG over time. Unlike in the developmental context, no photoreceptor-like cells were formed. Of note, these *Rbpj*-deficient MG did not incorporate EdU (Fig. S9a–b), indicating that Notch inhibition unlocks a slow, inefficient transdifferentiation process without inducing cell cycle activation.

**Figure 2.**
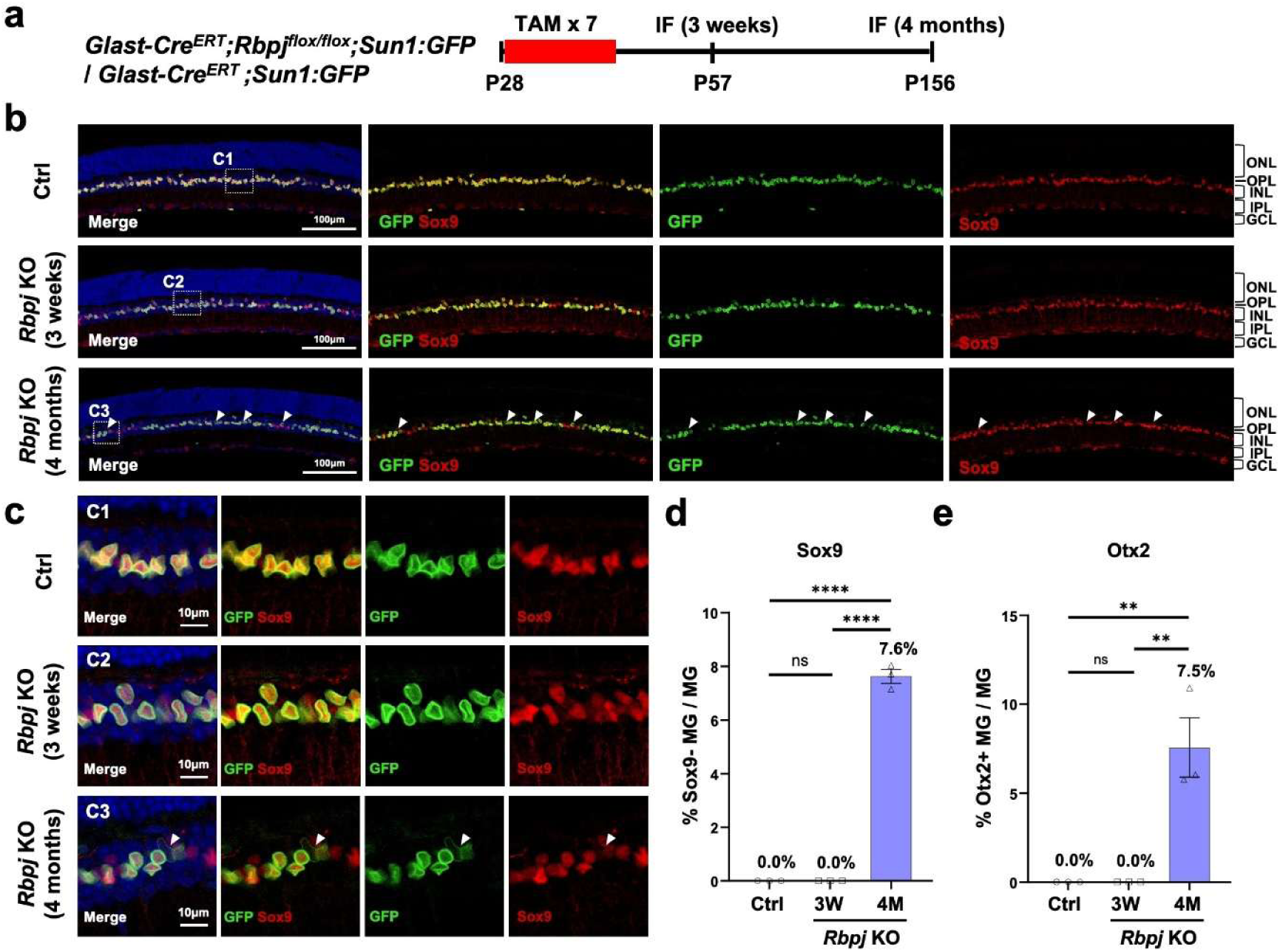
*Rbpj* deletion in adult MG induces limited dedifferentiation. (a) Schematic illustration of MG dedifferentiation and reprogramming experiment. (b) Representative immunostaining of Sox9 on retinal sections from *Glast-Cre^ERT^;Sun1:GFP* and *Glast-Cre^ERT^;Rbpj^flox/flox^;Sun1:GFP* mice in different timepoints post TAM injection. The white arrows refer to GFP+ Sox9- cells. (c) Magnified views of the highlighted regions in (b). (d) Percentage of GFP+ Sox9- cells in overall GFP+ cells. n=3 mice, data are presented as mean ± SEM. ns=not significant, \*\*\*\**P* < 0.0001, by one-way ANOVA with Tukey’s post hoc test. (e) Percentage of GFP+ Otx2+ cells in overall GFP+ cells. n=3 mice, data are presented as mean ± SEM. ns=not significant, \*\**P* < 0.01, by one-way ANOVA with Tukey’s post hoc test.

### Notch inhibition reduces but does not abolish CCA-induced MG proliferation

Since direct conversion depletes MG, we aimed to combine CCA-induced proliferation to expand the MG pool with *Rbpj* KO-driven reprogramming. Because Notch inactivation drives premature cell-cycle exit during development(*31–33*), we first determined if *Rbpj* deletion would antagonize the mitogenic activity of CCA. We administrated TAM from P28 to P35 to induce MG labeling and *Rbpj* knockout, followed by CCA injection at P36 (Fig. S11a). Subsequently, EdU was administrated intraperitoneally daily for 21 days, spanning the major time window of MG proliferation (Fig. S10a-c), and the mice were harvested to assess MG proliferation rates (Fig. S11a).

Remarkably, even in the absence of Notch signaling, CCA maintained a robust capacity to drive MG proliferation. Although the total number of EdU+ MG was moderately decreased in *Rbpj*-/- mice compared to wild type and *Rbpj*+/- controls, a substantial population of MG still successfully proliferated (Fig. S11b-c). Reversing the orders of TAM and CCA treatments yielded similar robust results (Fig. S11d-g). While this moderate reduction could reflect a minor anti-proliferative effect of Notch inhibition, it is also possible that *Rbpj* deletion down-regulates GFAP promoter activity in dedifferentiated MG and thereby lowers vector expression. Crucially, the persistence of numerous proliferated cells demonstrates that *cyclin D1* overexpression and *p27^Kip1^* suppression overrode these factors. This confirms that these downstream cell-cycle regulators act as potent drivers of proliferation independent of upstream Notch signaling (*34*).

### Combining *Rbpj* deletion and CCA treatment induce robust MG dedifferentiation and ectopic expression of Otx2

We next evaluated the reprogramming efficiency of the combined treatment of CCA and *Rbpj* KO in *Glast-Cre^ERT^;Rbpj^flox/flox^;tdT or Sun1:GFP* mice. Ideally, *Rbpj* deletion should be induced immediately after MG proliferation. However, CCA-driven MG proliferation in adult mice is not a synchronized process, with most MG proliferations occurring from week 1 to week 6 after CCA injection (Fig. S10a-c). To ensure precise lineage tracing and minimize confounding effects of CCA on gene expression, we adopted the protocol of administrating TAM immediately prior to CCA injection (Fig. S12a). At 3 weeks post-treatment, MG in all groups retained Sox9 expression (Fig. S12b-d). By 4 months, approximately 27.8% of GFP-positive MG-derived cells in the combination group had lost Sox9 expression, a significantly higher fraction than in the CCA alone (1.5%) or *Rbpj* KO alone (7.6%) groups (Fig. S12b-e). Importantly, the total number of Sox9+ MG remained higher in the combination group than that in the control, indicating that the combined treatment did not deplete the glial pool (Fig. S12f).

The enhanced dedifferentiation was accompanied by progressively more robust neurogenesis over time. Lineage tracing revealed no Otx2⁺ MG-derived cells in the combination group at 3 weeks post-treatment (Fig. 3a–d). By 2 months, approximately 9.2% of MG-derived cells had acquired Otx2 expression, and this proportion continued to rise to 27.4% by 4 month (Fig. 3b–e). This progressive increase indicates that MG-to-neuron conversion in the combination group unfolds gradually over months rather than as an abrupt switch. Compared with the CCA-alone and *Rbpj* deletion-alone groups, the percentage of Otx2⁺ MG-derived cells in the combined treatment group increased by approximately 20-fold and 3-fold, respectively (Fig. 3e; Fig. S13a). Injection of a control AAV_7m8_-GFAP-GFP vector did not increase the percentage of Otx2+ MG (Fig. S14a-c), indicating that the enhanced MG reprogramming was driven by the transgenes rather than by the AAV vector itself or the injection procedure.

**Figure 3.**
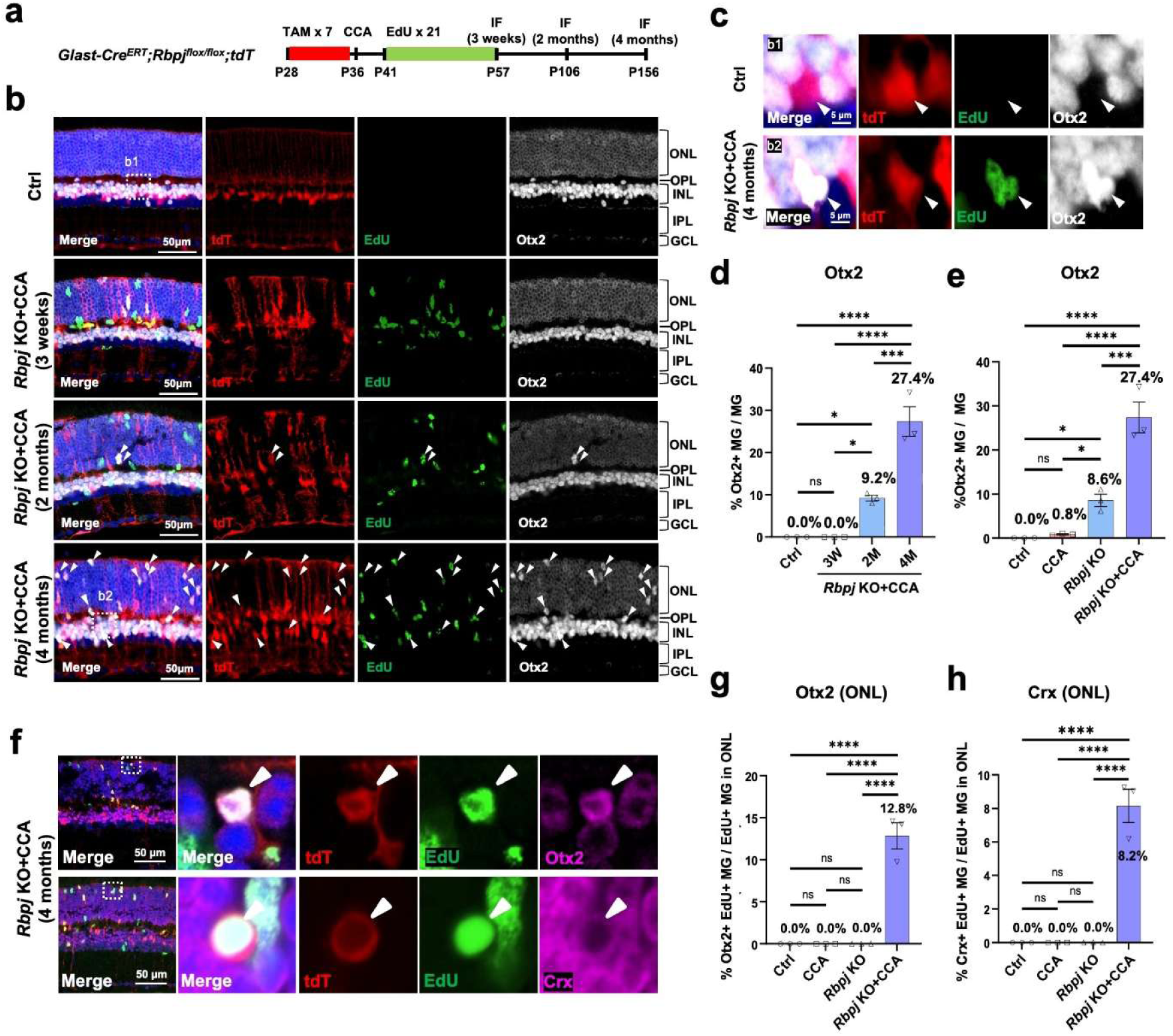
*Rbpj* KO and CCA synergistically increase Otx2+ cell formation from MG. (a) Schematic illustration of the neurogenesis assessment experiment. (b) Representative immunostaining of EdU and Otx2 on retinal sections. The white arrows refer to tdT+ Otx2+ cells. (c) Magnified views of the highlighted regions in (b). The white arrows refer to tdT+ Otx2+ cells. (d) Percentage of tdT+ Otx2+ cells in overall tdT+ cells. 3W: 3 weeks, 2M: 2 months, 4M: 4 months, n=3 mice, data are presented as mean ± SEM. ns=not significant, \*\*\**P* < 0.001, by one-way ANOVA with Tukey’s post hoc test. (e) Percentage of tdT+ Otx2+ cells in overall tdT+ cells. n=3 mice, data are presented as mean ± SEM. ns=not significant, \**P* < 0.05, \*\*\**P* < 0.001, \*\*\*\**P* < 0.0001, by one-way ANOVA with Tukey’s post hoc test. (f) Representative immunostaining of EdU and Otx2 or Crx on retinal sections. (g) Percentage of tdT+ EdU+ Otx2+ cells in ONL tdT+ EdU+ cells. N=3 mice, data are presented as mean ± SEM. Ns=not significant, \*\*\*\**P* < 0.0001, by one-way ANOVA with Tukey’s post hoc test. (h) Percentage of tdT+ EdU+ Crx+ cells in ONL tdT+ EdU+ cells. N=3 mice, data are presented as mean ± SEM. Ns=not significant, \*\*\*\**P* < 0.0001, by one-way ANOVA with Tukey’s post hoc test.

Given that photoreceptor loss is a leading cause of incurable blindness worldwide(*35*, *36*), reprogramming MG into photoreceptors represents a promising therapeutic strategy. We examined whether MG-derived cells adopted photoreceptor identities. In the combination group, a subset of postmitotic MG in the ONL upregulated photoreceptor markers, including Otx2 (12.8% of ONL tdT+ EdU+ MG) and Crx (8.2% of ONL tdT+ EdU+ MG), consistent with potential progression toward photoreceptor fate (Fig. 3f–h). The soma of some MG-derived cells in the ONL also adopted a photoreceptor cell-like circular morphology but without any structure of outer segment (Fig. 3f). However, only about 2% of ONL MG-derived cells expressed *Nrl*, an important factor regulating the rod photoreceptor specification(*37*) (Fig. S13b-c), which may limit maturation into fully developed rod photoreceptors.

In summary, these findings collectively demonstrate that the combined treatment of CCA and *Rbpj* deletion significantly enhances both the dedifferentiation of MG and the subsequent generation of retinal neurons compared to either treatment alone.

### *Rbpj* deletion promotes the formation of neuronal progenitor cells from MG

To dissect the transcriptional trajectory of reprogramming, we performed snRNA-seq on purified MG from *Glast-Cre^ERT^;Rbpj^flox/flox^;Sun1:GFP* mice (Fig. 4a). We analyzed four experimental groups, the untreated control (Ctrl), CCA treatment (CCA), *Rbpj* deletion (*Rbpj* KO), and combined *Rbpj* deletion and CCA treatment (*Rbpj* KO+CCA), at 1 week, 3 weeks, and 4 months post-treatment (Fig. 4a).

**Figure 4.**
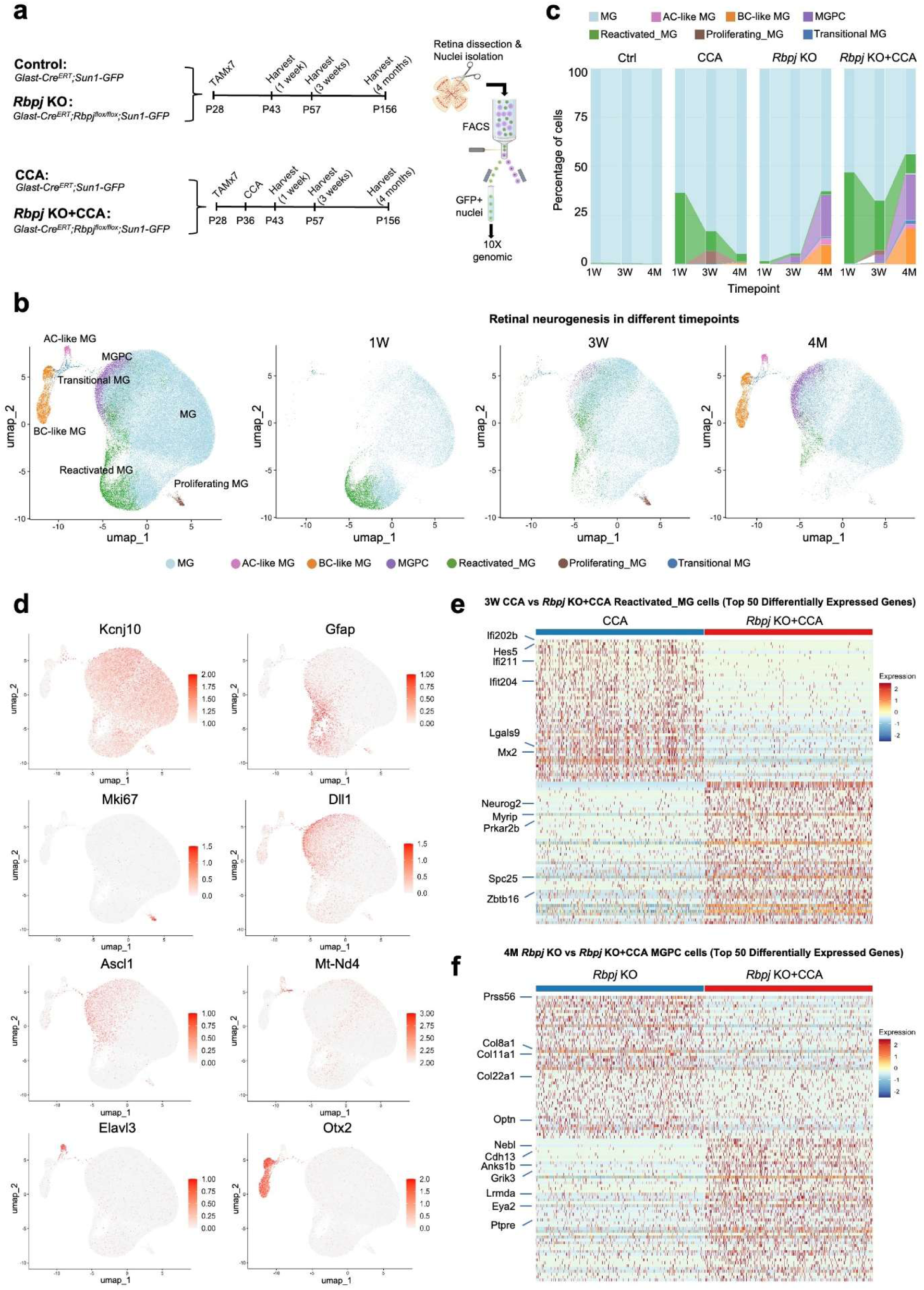
snRNA-seq analysis shows that *Rbpj* KO and CCA promote MG reprogramming. (a) Schematic illustration of the snRNA-seq experiment. To induce *Rbpj* deletion and GFP expression in MG, we administered TAM for 7 consecutive days in *Glast-Cre^ERT^;Sun1:GFP* and *Glast-Cre^ERT^;Rbpj^flox/flox^;Sun1:GFP* mice from P28. At P36, CCA was injected into the CCA and *Rbpj* KO+CCA groups, while control and *Rbpj* KO mice were uninjected by any AAV vector. 3-4 retinas from mice at indicated ages were pooled together for nuclei extraction, and MG nuclei were then isolated by GFP signal via FACS for snRNA-seq. (b) UMAP plot of snRNA-seq data from four treatment groups (Ctrl, CCA, *Rbpj* KO, *Rbpj* KO+CCA) at different timepoints, and separated UMAP based on different timepoints. clusters were identified based on known marker gene expression. (c) Proportions of cell clusters within Ctrl, CCA, *Rbpj* KO and + groups at different timepoints. (d) Feature plots highlighting the cluster of quiescent MG (*Kcnj10*), proliferating MG (*Mki67*), reactivated MG (*Gfap*), MGPC (*Dll1*, *Ascl1*), transitional MG (*Mt-Nd4*), AC-like MG (*Elavl3*), and BC-like MG (*Otx2*). (e) Heatmap showing the expression of top 50 differentially expressed genes (DEGs) of 3-week reactivated MG between CCA and *Rbpj* KO+CCA (p<0.05). (f) Heatmap showing the expression of top 50 DEGs of 4-month MGPC between *Rbpj* KO and *Rbpj* KO+CCA (p<0.05).

No significant batch effects were observed across the treatment groups (Fig. S15a). Following quality control filtering and the exclusion of native mature neurons, which were uniformly distributed across all three timepoints and treatment groups, approximately 3,500 to 12,000 cells remained for downstream snRNA sequencing analysis (Fig. S15a-b). The uniform manifold approximation and projection (UMAP) visualization of snRNA-seq data revealed dynamic cell cluster transitions over time (Fig. 4b-c). Clustering analysis identified eight MG-related populations: quiescent MG, reactivated MG, proliferating MG, MGPCs, transitional MG, amacrine cell-like MG and bipolar cell-like MG, annotated using known retinal markers (Fig. 4b–d, Fig. S15c-d, Fig. S16a-b).

At 1 week, prior to peak transgene expression, MG in the control and *Rbpj* KO groups remained quiescent, expressing high levels of MG genes (*Aqp4*, *Rlbp1*, *Kcnj10*) (Fig. 4b–d, Fig. S16a–c, Fig. S17a). In contrast, CCA treatment drove cells into a “reactivated” state marked by upregulation of the gliosis genes (*Gfap*, *Vim*), likely a response to AAV infection (Fig. 4b–d, Fig. S16a–c, Fig. S17a). At this time, no proliferating cells were observed in the CCA-injected eyes, as the AAV-mediated transgene expression had not reached the level required to drive the cell cycle.

At 3 weeks, the divergence between treatments became evident. In the CCA treated groups, MG (6.4% in CCA and 2.2% in *Rbpj* KO+CCA) proliferated (Fig. S16c). A subset of MG in *Rbpj* KO (3.4%) and *Rbpj* KO+CCA (4.0%) upregulated neurogenic factors (*Neurog2*, *Ascl1*, *Dll1*), marking them as MGPCs (Fig. 4b–d, Fig. S16a–c, Fig. S17a). A small fraction (<1%) of nascent AC-like cells (expressing *Elavl3*, *Rbfox3*, *Caln1*) and BC-like cells (expressing *Gsg1*, *Pcdh17*, *Lgr5*) emerged in *Rbpj* KO MG at this stage (Fig. 4b–d, Fig. S16a–c, Fig. S17a). By 4 months, while CCA-only MG had largely reverted to quiescence with minimal neurogenesis, the *Rbpj* KO and *Rbpj* KO+CCA groups sustained high percentages of MGPCs (21.5% and 23.5%, respectively) (Fig. 4b–d, Fig.S16b–c, Fig. S17a), confirming that Notch inhibition drives MGPC formation. Neurogenesis progressed significantly by this stage, with the *Rbpj* KO alone and *Rbpj* KO+CCA treatments yielding a higher percentage of AC-like cells (*Rbpj* KO, 3.2%; *Rbpj* KO+CCA, 2.2%) and BC-like cells (*Rbpj* KO, 9.9%; *Rbpj* KO+CCA, 18.3%) compared to the CCA-only group (Fig. 4b–d, Fig. S16b–c, Fig. S17a), Additionally, a unique “transitional” cluster enriched for mitochondrial gene (*mt-Cytb*, *mt-Nd4*) appeared in the *Rbpj* KO+CCA group (Fig. 4b–d, Fig. S16b–c, Fig. S17a), suggesting a metabolically active state conducive to reprogramming.

To elucidate the mechanism by which *Rbpj* deficiency promotes neurogenic potential, we performed differential gene expression analysis on 3-week reactivated MG, a critical inflection point between quiescence and commitment to MGPC. At this time point, the CCA-treated reactivated MG cluster exhibited heightened expression of genes associated with quiescent MG (*Hes5*, *Lgals9*) (Fig. 4e). The elevated expression levels of genes linked to quiescent MG may potentially guide reactivated MG back to a quiescent state (Fig. 4e). In contrast, the *Rbpj* KO+CCA-treated reactivated MG cluster demonstrated extensive expression of neurogenic genes, including *Neurog2*, *Zbtb16*, *Myrip*, and *Prkar2b*. (Fig. 4e). *Neurog2* and *Zbtb16* are well-established downstream genes regulated indirectly by the Notch–Rbpj pathway through the pro-neural differentiation programs(*38–40*). Their upregulation therefore reflects secondary effects of *Rbpj* loss and supports enhanced neurogenic reprogramming in the *Rbpj* KO+CCA group. Interestingly, in *Rbpj* KO+CCA-treated reactivated MG clusters, there was a significant suppression of genes associated with the interferon (IFN) pathway, including *Ifi211*, *Ifit202b*, and *Mx2* (Fig. 4e). As IFN signaling is known to play a crucial role in regulating the plasticity of MG(*41–43*), its downregulation, coupled with the activation of neurogenic factors, likely catalyzed the formation of MGPCs.

DEG analysis of 4-month MGPC populations revealed distinct transcriptional programs between the *Rbpj* KO and *Rbpj* KO+CCA groups (Fig. 4f). The *Rbpj* KO+CCA MGPCs upregulated genes associated with early retinal progenitor maintenance (*Nebl*, *Eya2*) and retinal development (*Lrmda*, *Ptpre*, *Grik3*) (Fig. 4f). Conversely, the *Rbpj* KO MGPCs preferentially expressed gliosis-related genes, such as *Prss56*, *Optn*, *Col8a1*, *Col22a1*, and *Col11a1* (Fig. 4f)(*44–46*). Given that retinal gliosis impedes MG reprogramming in mammals(*14*), these data support the idea that combining *Rbpj* deletion with CCA enhances regenerative potential by boosting early progenitor programs while mitigating the gliotic responses.

### CCA promotes BC subtype differentiation of the *Rbpj*-deficient MG

To investigate the heterogeneity of the newly generated neuron, we performed sub-clustering analysis focused specifically on the BC-like cells, which constituted a major reprogrammed neuron-like cluster. Using established BC subtype markers(*47*), we identified three distinct BC subtypes: OFF-cone BCs (identified by *Pcdh17*, *Zfhx4*, *Lrrtm3*, *Chrm2*), ON-cone BCs (identified by *Isl1*, *Pde8a*, *Esrrg*, *Ryr3*, *Cdh9*, *Grm6*), and rod BCs (identified by *Isl1*, *Prkca*, *Cep112*, *Lrrtm4*) (Fig. 5a-c). Comparative analysis revealed that *Rbpj* deletion and CCA generated a higher abundance of these BC subtypes than either CCA or *Rbpj* deletion treatment alone (Fig. 5d).

**Figure 5.**
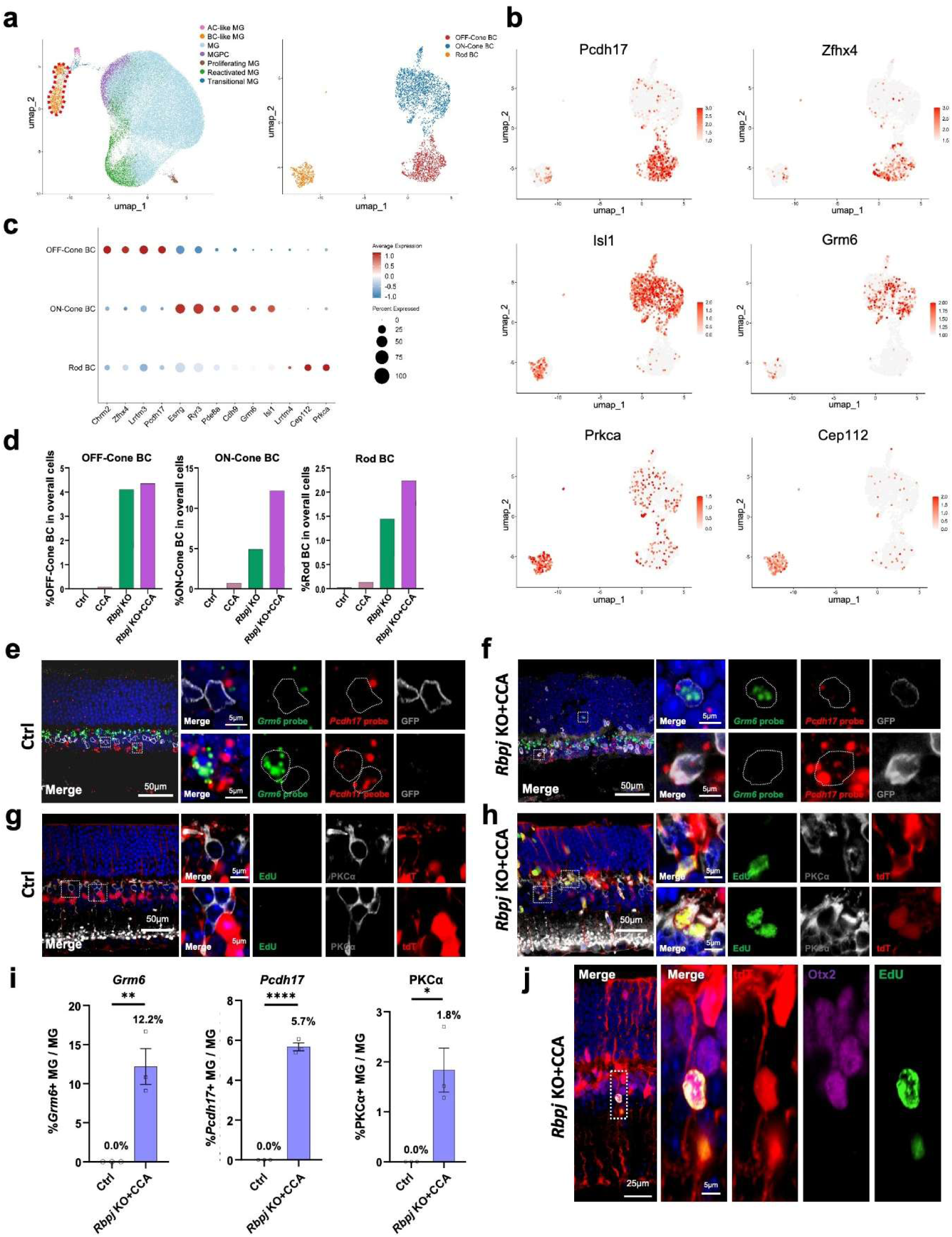
CCA and *Rbpj* deletion drive BC subtype differentiation. (a) Subclusters of the BC-like population. The BC-like MG (outlined in red) was used for subclustering analysis. (b) Feature plot of BC-like subtypes showing the cluster of OFF-cone BC (*Pcdh17*, *Zfhx4*), ON-cone BC (*Isl1*, *Grm6*) and rod BC (*Isl1*, *Prkca*, *Cep112*). (c) Dot plot showing gene expression and cell percentages for OFF-cone BC, ON-cone BC and rod BC. (d) Percentage of BC subtypes in different treatment groups. (e-f) *Pcdh17* and *Grm6* mRNA *in situ* hybridization in the Ctrl and *Rbpj* KO+OCCA groups. The white dashed boxes indicate the position of the enlarged images. (g-h) Representative immunostaining of EdU and PKCα on retinal sections of the Ctrl and *Rbpj* KO+CCA samples. The white dashed boxes indicate the position of the enlarged images. (i) Percentage showing GFP+ *Grm6*+ cells in overall GFP+ cells, GFP+ *Pcdh17*+ cells in overall GFP+ cells and GFP+ PKCα+ cells in overall GFP+ cells in the Ctrl and *Rbpj* KO+CCA groups. N=3 mice, \**P* < 0.05, \*\**P* < 0.01, \*\*\*\**P* < 0.0001, by unpaired two-tailed student’s t-test. (j) Representative immunostaining of MG-derived cells with bipolar cell morphology.

To validate the snRNA-seq findings, we performed RNA *in situ* hybridization on *Pcdh17*, *Grm6*, and immunostaining on PKCα to identify OFF-cone BCs, ON-cone BCs, and rod BCs, respectively (Fig. 5e-h). No MG in the control group co-localized with any of these BC markers. The proportions of newborn BC subtypes among MG-derived cells, as quantified by staining, were similar to the proportions observed in the snRNA-seq data (Fig. 5i). Furthermore, some MG-derived neuron-like cells, which are EdU+ Otx2+ tdT+, exhibited morphological changes towards BC, characterized by the retraction of their apical glial processes and the adoption of BC nucleus shapes (Fig. 5j). Collectively, these results indicate that *Rbpj* KO+CCA promoted the generation and maturation of BC subtypes formation.

To better understand the trajectory of neuronal regeneration, we performed pseudotime analysis (Fig. S18a). This revealed a clear and sequential neurogenic order, progressing from resting MG to neurogenic MGPCs, and finally to BC-like and AC-like cells. This progression was closely associated with the downregulation of MG-specific genes (e.g., *Hes5*, *Kcnj10*, *Slc1a2*) and the upregulation of neurogenic factors (e.g., *Neurog2*, *Dll1*, *Eya2*) (Fig. S18b-d). The *Rbpj* KO+CCA treatment group exhibited a more rapid shift in the gene-to-cell ratio trend line towards the BC formation branch compared to the CCA-alone and *Rbpj* KO-alone groups (Fig. S18e), which may explain the increased diversity and abundance of BC-like cells in the *Rbpj* KO+CCA group.

We also investigated whether AC subtypes formed from the MG. Although this cluster could be separated into two distinct sub-clusters, they did not clearly correspond to specific AC subtypes using known marker(*48*), such as *Chat* for Starburst ACs, *Dab1* for A17 ACs, and *Slc6a9* for nGnG ACs (Fig. S19a). Immunostaining for HuC/D confirmed the formation of AC-like cells, consistent with the snRNA-seq detection (Fig. S19b-c). Quantification further confirmed that the loss of *Rbpj* promotes AC formation compared to the CCA-alone treatment group (Fig. S19d-e).

While the neuron-like clusters were best classified as BC-like and AC-like based on their distinct marker gene expression, they also exhibited mixed expression of genes associated with other retinal neuronal types, including *Tubb3*, *Myt1l*, and *Grin1, which are the signature genes of RGC,* and *Crx*, *Prom1*, *Epha10*, *Gucy2e*, *Scg3* which are signature genes associated with photoreceptor lineages (Fig. S20a), suggesting that the regenerated cells exist in a hybrid state. Notably, we did not detect *Nrl* expression in the newborn neuron clusters, which is essential for rod photoreceptor specification(*37*) (Fig. S20a). The discrepancy in Nrl detection between snRNA-seq and prior immunofluorescence staining suggests that Nrl+ cell generation is stochastic and occurs at low frequency (Fig. S13b–c, Fig. S20a). Moreover, relative to the *Rbpj* KO group, neuron-like clusters in the *Rbpj* KO+CCA group showed upregulated expression of genes typically enriched in both RGCs and photoreceptors, such as *Tmem132d*, *Cntn5*, *Ryr2*, *Mef2c*, *Meis2*, and *Cngb1* (Fig. S20b).

Together, these mixed lineage signatures indicate that the newly generated neurons remain in a plastic, incompletely committed state and likely require additional cues, such as Nrl, to achieve full maturation and terminal specification. Moreover, combining *Rbpj* deficiency with CCA treatment enhances the acquisition of abundant and distinct neuronal characteristics in the newborn neurons.

### Increased chromatin accessibility of neurogenic factors underlies CCA-induced neuronal formation from MGPCs

We next sought to elucidate how CCA promotes neurogenesis. We hypothesized that CCA-induced MG proliferation remodels the chromatin landscape to facilitate reprogramming, a mechanism analogous to those observed in somatic cell reprogramming and cardiac regeneration(*49*, *50*). To test this hypothesis, we profiled chromatin accessibility using snATAC-seq on MG nuclei purified from four groups at 4 months post-treatment, including the untreated control, CCA, *Rbpj* KO, and the combined *Rbpj* KO+CCA group (Fig. 6a).

**Figure 6.**
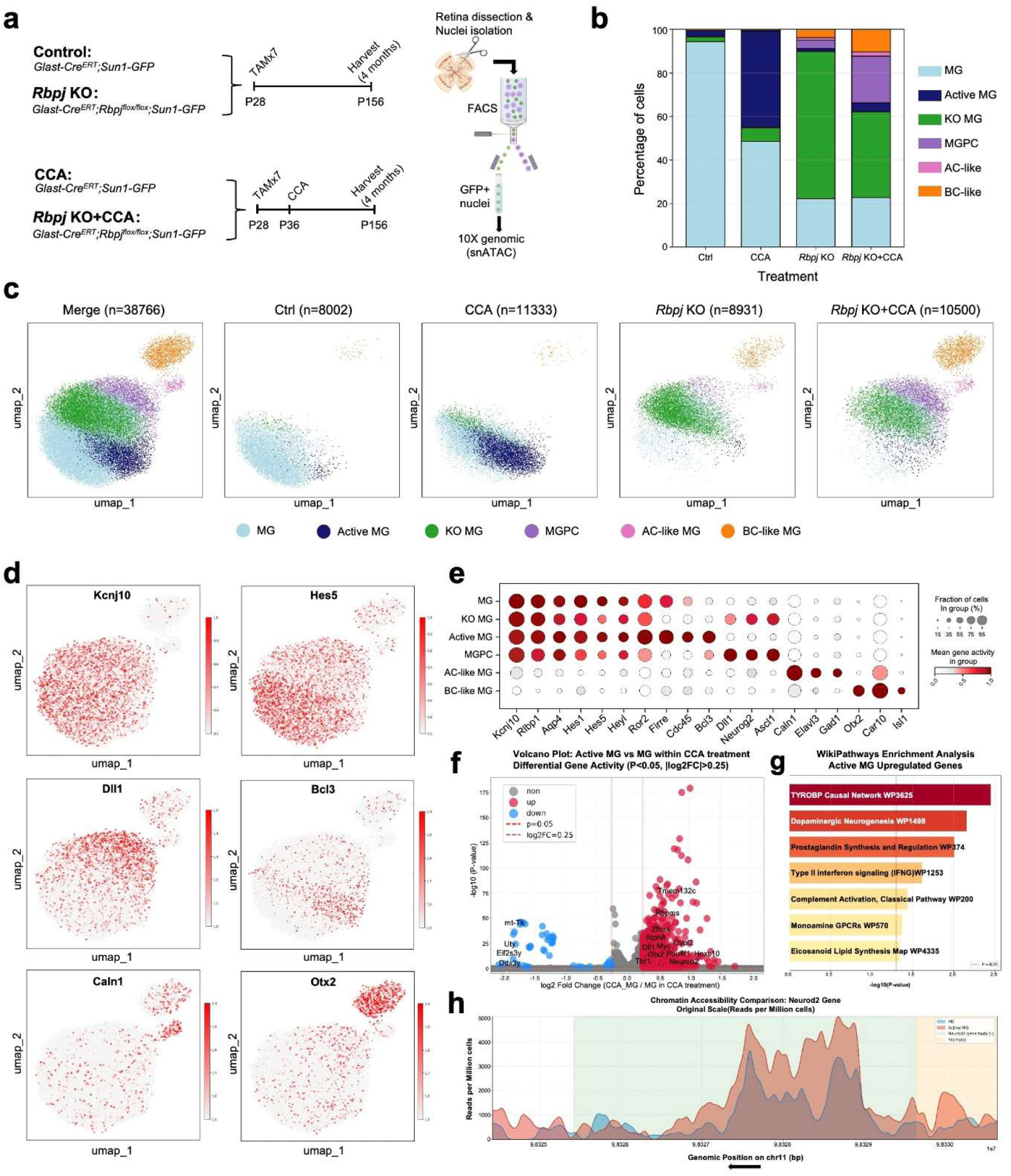
snATAC-seq analysis suggests increased chromatin accessibility of the neurogenic genes by CCA. (a) Schematic illustration of the snATAC-seq experiment. (b) Proportions of cell clusters within Ctrl, CCA, *Rbpj* KO and *Rbpj* KO+CCA groups. (c) UMAP plot of snATAC-seq data from four treatment groups (Ctrl, CCA, *Rbpj* KO, *Rbpj* KO+CCA). (d) Feature plots highlighting the cluster of quiescent MG (*Kcnj10*), KO MG (*Hes5*), active MG (*Bal3*), MGPC (*Dll1*), AC-like MG (*Caln1*), BC-like MG (*Otx2*). (e) Dot plot showing gene activity and cell percentages for quiescent MG, KO MG, active MG, MGPC, AC-like MG, BC-like MG. (f) Volcano plot showing the differential gene activity of MG and active MG within CCA treatment group (p<0.05). Red dots indicate the genes upregulated in active MG and blue dots indicate genes downregulated in active MG. (g) WikiPathways enrichment analysis of upregulated genes in active MG (p<0.05). (h) Increased chromatin accessibility at the Neurod2 locus was observed in the active MG. The black arrow shows the transcription direction.

No batch effects were detected across treatments, and clusters were annotated using canonical cell type-specific markers (Fig. S21a–c). After quality control and removal of native mature neurons, which uniformly represented across groups, approximately 8,000–11,000 cells remained for downstream analysis (Fig. 6b–c). Clustering resolved six cell types based on marker accessibility: resting MG, active MG, *Rbpj* KO MG (KO MG), MGPC, AC-like, and BC-like, with distinct compositional shifts across treatments (Fig. 6b–c). As expected, MG in the control group exhibited a quiescent chromatin state, with high accessibility at classic MG marker genes (e.g., *Kcnj10*, *Rlbp1*, *Aqp4*) (Fig. 6d-e). A unique active MG cluster, showing markedly increased accessibility at loci involved in cell cycle regulation (e.g., *Firre*, *Cdc45*, *Bcl3*), was characterized in the CCA treated group (Fig. 6d-e). In *Rbpj* deficient MG, accessibility at Notch pathway effectors (*Hes1*, *Hes5*, *Heyl*) was markedly reduced relative to resting MG (Fig. 6d–e). Notably, the *Rbpj* KO+CCA combination produced more pronounced chromatin changes associated with MGPC, AC-like, and BC-like states compared to *Rbpj* KO alone, underscoring the contribution of CCA to chromatin remodeling (Fig. 6d–e).

A direct comparison of resting MG versus active MG within the CCA treatment group revealed broad increases in chromatin accessibility (Fig. 6f), with upregulated loci linked to retinal development and neurogenesis, including *Neurod2*, *Dll1*, and *Otx2* (Fig. 6f). WikiPathways enrichment indicated strong associations with dopaminergic neurogenesis (Fig. 6g). Genome browser tracks further showed elevated accessibility across the *Neurod2* promoter and gene body in active MG relative to resting MG, consistent with the activation of this key neurogenic regulator (Fig. 6h).

In summary, our snATAC-seq data support a model wherein CCA-induced proliferation remodels the chromatin landscape of MG, priming them for reprogramming by opening key neurogenic loci.

### MG-derived daughter cells exhibit long-term survival

To evaluate long-term outcomes of MG reprogramming, a cohort of mice treated with *Rbpj* KO and CCA were aged to 9 months post-treatment (Fig. 7a). At this late time point, MG-derived cells, marked by the tdT reporter, displayed a more regular, circular nuclear morphology reminiscent of mature retinal neurons, though fully mature rod photoreceptors were not detected (Fig. 7b). Quantification showed an upward trend in the proportion of Otx2+ tdT+ cells among total tdT+ cells, consistent with a gradual, ongoing MG-to-neuron reprogramming process (Fig. 7b-c). However, both the percentage of Otx2+ EdU+ tdT+ cells within the EdU+ tdT+ population and the absolute number of EdU+ cells declined slightly over time (Fig. 7d–e). This data suggests that while most MG-derived daughter cells persist long-term, a small subset of proliferated MG-derived neurons may ultimately undergo apoptosis.

**Figure 7.**
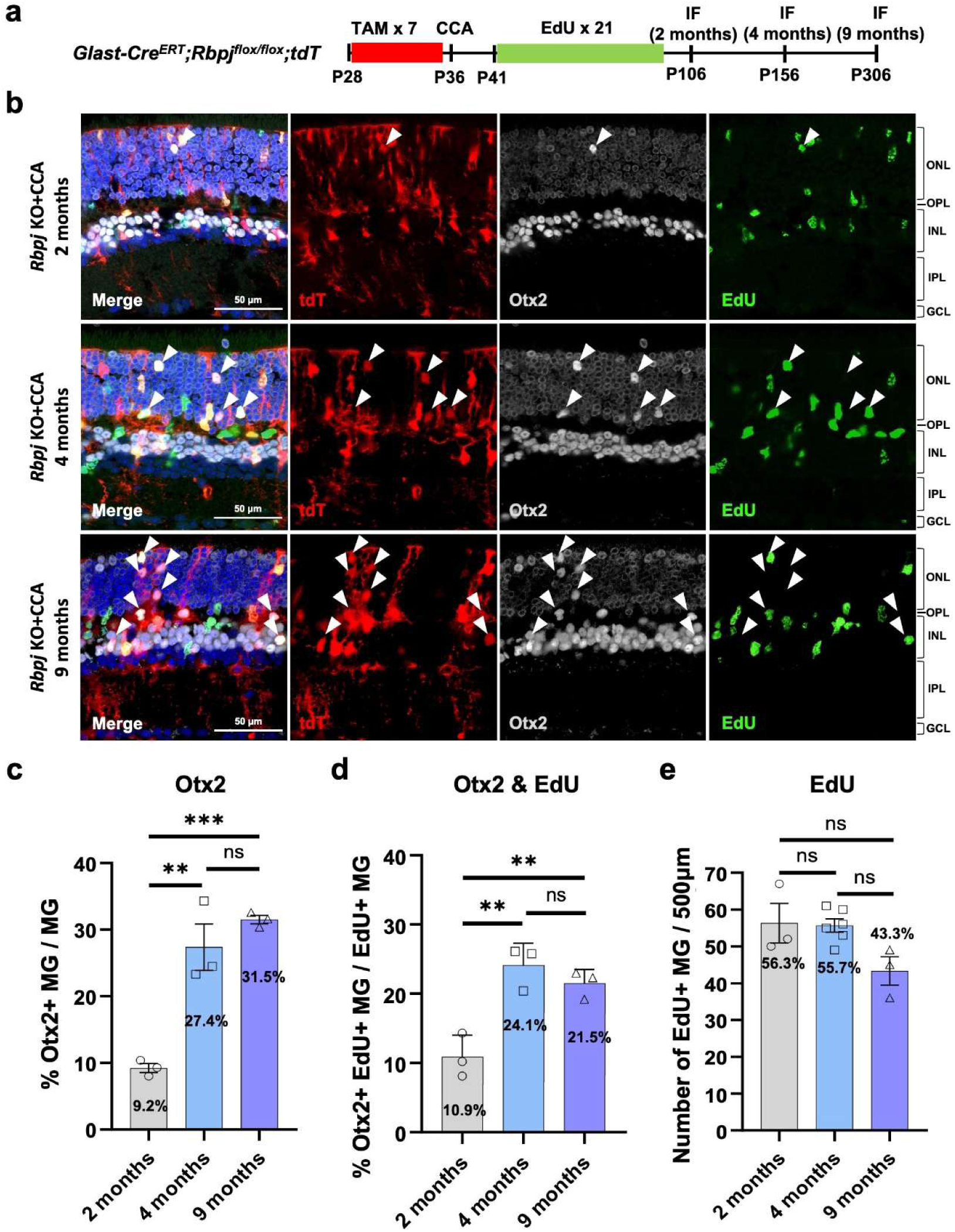
Neurons derived from MG exhibited long-term survival capabilities. (a) Schematic illustration of the experiment assessing long-term cell survival. (b) Representative immunostaining of EdU and Otx2 on retinal sections. The white arrows refer to tdT+ Otx2+ cells. (c) Percentage of tdT+ Otx2+ cells in overall tdT+ cells, n=3 mice, data are presented as mean ± SEM. ns=not significant, \*\**P* < 0.01, \*\*\**P* < 0.001, by one-way ANOVA with Tukey’s post hoc test. (d) Percentage of tdT+ EdU+ Otx2+ cells in overall tdT+ EdU+ cells, ns=not significant, \*\**P* < 0.01, by one-way ANOVA with Tukey’s post hoc test. (e) Number of EdU+ cells per 500 µm, ns=not significant, by one-way ANOVA with Tukey’s post hoc test.

To evaluate retinal structure following treatment, we performed ZO1 staining to visualize the outer limiting membrane (OLM) (Fig. S22a). The OLM integrity in the *Rbpj* KO+CCA group at 9 months was comparable to that of untreated controls, suggesting that sufficient MG persist to support junctional complexes (Fig. S22b). Next, to assess whether the combined *Rbpj* KO and CCA treatment compromises retinal function, we performed structural and functional analyses at 4 months post-treatment (Fig. S23a). Optical coherence tomography (OCT) revealed that retinal layer organization and ONL thickness were comparable among the uninjected eyes, GFP AAV-injected control eyes, and CCA-treated eyes, demonstrating that overall retinal architecture was well preserved (Fig. S23b–c). Consistently, optomotor response testing revealed no significant differences in visual acuity across groups, suggesting that visual function remained intact (Fig. S23d). Furthermore, electroretinography (ERG) demonstrated that scotopic and photopic a- and b-wave amplitudes were unaffected by the treatment, confirming that light responses from photoreceptor and inner retinal neuron were preserved (Fig. S23e–h). Taken together, these findings demonstrate that combined *Rbpj* KO and CCA treatment achieves neurogenesis without compromising retinal structure and functional visual circuitry.

In summary, our findings demonstrate that the majority of MG-derived cells possess inherent long-term stability, and that maintaining an adequate MG population is a prerequisite for achieving stable and functional regeneration.

## Discussion

Retinal degeneration is characterized by the progressive loss of retinal neurons, leading to irreversible vision impairment. In teleost fish, retinal injury triggers functional regeneration, with MG serving as the source of regenerated neurons. In contrast, adult mammalian retinas lack regenerative capacity because MG fail to re-enter the cell cycle, dedifferentiate, and subsequently differentiate into neurons. In our previous study, we showed that simultaneous knockdown of *p27^Kip1^* and overexpression of *cyclin D1* effectively drives mouse MG to re-enter the cell cycle in the absence of injury. However, most postmitotic MG underwent only transient dedifferentiation, and only a small fraction (∼1%) differentiated into neuron-like cells(*10*). In the present study, scRNA-seq and RNA *in situ* hybridization revealed persistent activation of Notch signaling in postmitotic MG, which restricts their reprogramming potential. Conditional deletion of *Rbpj*, the core transcriptional effector of Notch signaling, in late-stage RPCs promoted ectopic rod photoreceptor formation at the expense of MG differentiation, whereas *Rbpj* deletion in mature MG alone produced only limited neurogenic effects, suggesting that epigenetic landscape changes may contribute to more limited reprogramming potential from late RPC-to-MG transition. Notably, combining *Rbpj* knockout with the proliferative stimulus CCA treatment markedly enhanced neurogenesis from postmitotic MG. Immunofluorescence showed that ∼26% of MG daughter cells expressed neuronal marker Otx2 and HuC/D. snRNA-seq confirmed these results and revealed a clear trajectory of MG-derived neurogenesis, in which cells transitioned from a glial state toward BC-like and AC-like identities. Subtype analysis identified three BC subtypes: OFF-cone BCs, ON-cone BCs, and rod BCs, among MG-derived BC-like cells, whereas AC-like cells could not be further classified. MG-derived neuron-like cells also expressed genes characteristic of RGCs and photoreceptors, indicating enhanced lineage promiscuity. This hybrid transcriptomic profile, together with the immature morphology of newborn neurons, suggests incomplete or inefficient reprogramming.

Mechanistically, snATAC-seq revealed that CCA treatment increases chromatin accessibility at pro-neurogenic loci following MG proliferation, likely facilitating the improved reprogramming efficiency of *Rbpj*-/- MG. Notably, the majority of reprogrammed neurons survived for up to 9 months, while the MG pool remained intact, underscoring their preserved function to support retinal homeostasis.

MG are essential for maintaining retinal structural and functional integrity. They contribute to the blood-retinal barrier, regulate extracellular ion and neurotransmitter homeostasis, provide metabolic support to neurons, and mediate inflammatory responses(*51*, *52*). Thus, preserving MG number and function during reprogramming interventions is critical, as extensive MG loss or dysfunction of MG could disrupt retinal homeostasis and exacerbate degeneration(*17*). In zebrafish, MG regenerate neurons through asymmetric division, generating one self-renewing MG and one progenitor cell, thereby maintaining the glial pool(*53*, *54*). In our study, combined CCA treatment and *Rbpj* deletion effectively stimulated MG proliferation and reprogramming in the adult mouse retina without overt MG depletion. Because both daughter cells lacked *Rbpj*, a classic asymmetric division with one daughter have high Notch signaling and the other low is unlikely. Nonetheless, our strategy successfully regenerated neuron-like cells while maintained the MG pool, as more MG were generated to replace the reprogrammed MG. The long-term survival of newborn neurons and the stability of retinal architecture and function support the integrity of the MG pool. Although our strategy differs from zebrafish MG self-renewal, it still offers a viable regenerative approach that preserves glial homeostasis, a critical prerequisite for maintaining retinal homeostasis. Achieving a balance between neurogenesis and MG preservation represents a central challenge in glial-based regeneration. Excessive or widespread reprogramming risks depleting MG below a critical threshold or impairing their supportive functions. Future strategies should emphasize transient, partial, or spatially confined reprogramming to maintain a functional MG pool. For instance, replication-incompetent retroviral vectors(*55*, *56*) to knockout *Rbpj* in one daughter cell of a divided MG could ensure that the sibling cell remains a supportive glial cell, thereby maintaining retinal homeostasis while generating new neurons.

Multiple studies have explored the proliferative potential of MG in adult mice, including Wnt pathway activation, Hippo pathway inhibition, and forced manipulations of cell cycle regulators(*10–12*). However, unlike zebrafish, proliferating mouse MG rarely give rise to mature retinal neurons, regardless of the proliferation-inducing method. Our time-course snRNA-seq analysis of CCA-treated retinas confirmed this limitation: MG re-entered and exited the cell cycle but largely reverted to a quiescent glial state. What is the functional role of MG proliferation in retinal regeneration? Our findings imply that proliferation actively facilitates reprogramming rather than merely expanding MG numbers. Specifically, we demonstrate that CCA-induced proliferation triggers widespread chromatin opening at key neurogenic gene loci, including *Neurod2*, *Dll1*, and *Otx2*, in active MG compared to resting MG (Fig. 6f–h). This principle parallels findings in somatic cell reprogramming, where DNA replication and cell division facilitate the erasure of lineage-restrictive epigenetic marks to enhance reprogramming efficiency (*50*, *57–59*). Similarly, proliferative neural stem cells (NSCs) exhibit higher chromatin accessibility at pro-neural genes Otx2 and Ascl1, compared with aged NSCs that show reduced proliferative capacity(*60*, *61*). Enhancing NSCs proliferation with betacellulin upregulated Ascl1 and Nestin, thereby boosting neurogenesis in the olfactory bulb and dentate gyrus(*62*). Crucially, while this replication-coupled chromatin remodeling is insufficient to drive neurogenesis, it creates a permissive, epigenetically reset window. When combined with *Rbpj* deletion, which relieves Notch-mediated repression of downstream neurogenic factors such as *Ascl1*, *Neurog2*, *Otx2*, and *Neurod2*, to enable their effective occupancy and activation of the newly accessible chromatin. Taken together, our findings provide strong evidence that cell-division-mediated epigenetic resetting is conserved in MG and is essential for overcoming the barriers to retinal neurogenesis.

The Notch signaling pathway plays a pivotal role in regulating both cell proliferation and cell fate specification throughout retinal development(*31*, *32*). During development, Notch-mediated lateral inhibition ensures that diverse retinal cell types are generated in a spatially and temporally coordinated manner. Deletion of Notch1 or downstream effectors such as *Hes1* and *Rbpj* disrupts this balance, causing premature cell-cycle exit of retinal progenitor cells and increased production of early-born neurons, particularly photoreceptors, at the expense of later-born cell types, including MG(*31*, *63*). Conversely, forced expression of *Hes1* in progenitor cells drives gliogenesis and promotes MG differentiation(*64*). Align with this paradigm, *Rbpj* deletion during retinal development redirected the fate of late-stage progenitor cells, producing more mature rod photoreceptors at the expense of MG and BCs. In the mature retina, sustained Notch signaling maintains MG quiescence(*14*, *26*). Its conserved role across species has been well documented: inhibition of the Delta-Notch3-Hey1/Id2b pathway triggers zebrafish MG proliferation and regeneration(*18*, *65*, *66*). In the post hatch chick retina, Notch inhibition using a γ-secretase inhibitor likewise enhances retinal regeneration(*67*). In adult mice, simultaneous suppression of Notch signaling and deletion of Nfia/b/x synergistically promotes MG conversion into BCs and ACs(*25*). Similarly, virus-mediated Oct4 overexpression combined with Notch inhibition boosts the neurogenic competence of mammalian MG(*24*). Collectively, these findings position Notch inhibition as a central mechanism for unlocking MG plasticity. Together, these studies highlight Notch inhibition as a critical gatekeeper of MG plasticity. However, both our work and others indicate that Notch inhibition primarily generates BC- and AC-like neurons, with limited evidence for mature photoreceptor formation. This underscores an enduring barrier to producing the full repertoire of retinal neurons and suggests that additional lineage determinants, such as Nrl, may be required to guide MG toward photoreceptor fates.

Furthermore, we acknowledge that the precise morphological features, synaptic connectivity, and electrophysiological properties of MG-derived cells remain largely uncharacterized in the current study. As demonstrated by both our immunostaining and snRNA-seq data, the MG-derived neuron-like cells exhibit an incompletely mature transcriptomic profile with hybrid lineage signatures. While we observed initial morphological changes toward a bipolar cell identity, including retraction of apical glial processes and adoption of bipolar cell-like nuclear morphology (Fig. 5j), these cells lacked the fully elab orated dendritic and axonal arbors characteristic of mature BC. Moreover, we did not observe any MG-derived cells with photoreceptor outer segment or typical RGC morphology. Consequently, it is highly probable that functional synaptic connectivity and mature electrophysiological properties have not yet been established at the current stage of reprogramming. Future studies incorporating synaptic marker analysis, morphological reconstruction, and electrophysiological recordings will be essential to determine whether further optimization of our reprogramming strategy can drive these MG-derived neurons toward full functional maturation and successful circuit integration.

Regenerating retinal neurons from endogenous MG represents a promising frontier in regenerative medicine, yet the long-term survival of newly generated neurons is essential for clinical translation. Normally, mature retinal neurons depend on intrinsic survival mechanisms and extrinsic trophic support from the microenvironment, including trophic factors, such as brain-derived neurotrophic factor (BDNF), ciliary neurotrophic factor (CNTF), glial cell-line derived neurotrophic factor (GDNF) and nerve growth factor (NGF), synaptic integration for functional activity-dependent survival, and metabolic support from the RPE, and vasculature(*68*, *69*). In contrast, transplanted neurons often suffer low survival rate due to poor migration, inadequate support, limited synaptic integration, and immune rejection(*70–72*). For example, xenotransplantation of mouse induced pluripotent stem cell/mouse embryonic stem cell-derived RGCs exhibit only minimal medium-term survival in the host retina owing to immune barriers(*70–72*). While the survival of endogenous MG-derived neurons has been under explored, our study demonstrates a remarkably high long-term survival rate, with ∼80% of postmitotic MG-derived newborn neurons persisting for at least 9 months following combined *Rbpj* deletion and CCA treatment. We speculate that it may involve successful apical migration to their appropriate retinal layers (e.g., ONL and INL), where they can access layer-specific trophic support and integrate into local microenvironments. Their laminar positioning likely facilitates contact with presynaptic and postsynaptic partners, as well as access to survival signals from MG and photoreceptor cells, which are critical for preventing apoptosis. To further enhance the survival of newborn neurons from MG, future strategies should focus on recapitulating the endogenous survival mechanisms of the retina. This could involve co-expression of anti-apoptotic genes such as *Bcl-2* or *Xiap*(*73*, *74*) and supplemental trophic support (e.g., BDNF, CNTF), alongside synaptic integration through molecular guidance cues such as Semaphorins or Netrins(*75*, *76*). Regenerating neurons within a degenerating retinal environment, where vacated synaptic positions may facilitate new connections, may further enhance neuronal integration. The efforts to enhance the survival of regenerated neurons will be essential for advancing regenerative therapies.

Beyond survival, the appropriate laminar positioning of MG-derived neurons remains another challenge for retinal in vivo reprogramming. During retinal development, precise laminar positioning of neurons is guided by a coordinated interplay of cell-intrinsic transcriptional programs and extrinsic cues including cell adhesion molecules, guidance factors, and interactions with neighboring cells. In adult retina, many of these developmental cues are no longer present or active, which likely contributes to the failure of MG-derived neurons to migrate to their appropriate laminar positions. Interestingly, the vast majority of our divided MG cells remained localized within the ONL. Because the ONL is the physiological niche of photoreceptors, this preferential position could serve as an advantageous baseline layout for driving targeted photoreceptor differentiation in future work. In summary, our study establishes that combined *Rbpj* deletion and CCA-induced proliferation efficiently reprograms adult mouse MG into diverse retinal neurons that exhibit remarkable long-term survival, providing a robust platform for dissecting the molecular logic of mammalian retinal regeneration. Building upon this framework, future efforts to refine subtype specification, guide laminar positioning, and enhance synaptic integration will be critical to translating endogenous MG reprogramming into a viable therapeutic strategy for restoring vision in retinal degenerative diseases.

## Materials and Methods

### Animals

*Rbpj^flox/flox^* mice (strain: *Rbpj^em2Lutzy^*), *tdTomato* reporter mice (strain *B6.Cg-Gt(ROSA)26Sor^tm14(CAG-tdTomato)Hze^/J*)(*77*), *Sun1:GFP* reporter mice (Strain *B6.129-Gt(ROSA)26Sor^tm5.1(CAG-Sun1/sfGFP)Nat^/MmbeJ*)(*78*), and *Glast-Cre^ERT^* reporter mice (strain *Tg(Slc1a3-cre/ERT)1Nat/J*)(*79*) mice were purchased from the Jackson Laboratory. All mice were kept on a 12/12-hour light/dark cycle in the Laboratory Animal Research Unit, City University of Hong Kong. All animal procedures performed were approved by the Hong Kong Department of Health under Animals Ordinance Chapter 340 (Ref: (20–130) in DH/HT&A/8/2/5 Pt.2) and by the City University of Hong Kong Animal ethics committee (Ref: A-0264).

### AAV Production

AAV production was carried out following the previously described procedure (*80*). In brief, to produce the recombinant AAV_7M8_-GFAP-*cyclinD1*-*p27^Kip1^* shRNA-WPRE and AAV_7m8_-GFAP-GFP-WPRE, the HEK293T cells were transfected with a mixture of pAAV vector transgene plasmid, rep/cap packaging plasmid and adenoviral helper plasmid. At 96 hours post-transfection, both the culture medium and the transfected cells were harvested for AAV collection. The collected AAV underwent further purification through ultra-centrifugation in the iodixanol (OptiPrep) gradient at 147,000 x g at 4°C for 90 minutes. The iodixanol in the AAV solution was washed three times with PBS using Amicon 100K columns (EMD Millipore), and about 30µl of the final volume AAV was collected for downstream applications. The virus titration was determined through the protein SDS-PAGE method.

### Tamoxifen injection

To induce Cre recombinase expression in the majority of MG, tamoxifen (Sigma, dissolved in corn oil) was administered via intraperitoneal injection at a dosage of 50mg/kg.

### Intravitreal AAV injection

To stimulate MG proliferation, mice were intravitreally injected with AAV_7m8_-GFAP-*cyclinD1*-*p27^Kip1^* shRNA-WPRE and AAV_7m8_-GFAP-GFP-WPRE at a final concentration of 4E13vg/ml. Briefly, the eyelid was gently manipulated with tweezers to expose the eyeball. Subsequently, 1μl of the virus was precisely introduced into the intravitreal space using a custom angled glass pipette controlled by a FemtoJet (Eppendorf). The treatment was administered to the right eye of the animal, while the left eye remained untreated as a control.

### Retinal Cryosection and Immunohistochemistry

The mice were humanely euthanized using CO_2_ and cervical dislocation. Before enucleation, the eyeballs were marked ventrally to indicate the AAV injection site, and then the retinas were carefully dissected in PBS and fixed in 4% paraformaldehyde (PFA) at room temperature for 30 minutes. The fixed retinas underwent three-times washes with PBS (Product #10010023, Thermo Fisher Scientific) and were subsequently dehydrated in sequential sucrose solutions of 5%, 15%, and 30% for 15, 30, and 60 minutes, respectively. The retina was then immersed in a solution of optimal cutting temperature (OCT) and 30% sucrose in a 1:1 ratio at 4°C overnight before being embedded in cryomolds in a specific orientation for sectioning.

After freezing the tissue below -20°C, a series of 20μm sections were cut and mounted on glass slides using a cryostat machine (Thermo HM525NX Cryostat). For immunostaining, retinal sections were first blocked in 3% bovine serum albumin (BSA, #A9647, Sigma-Aldrich) and 0.1% Triton X-100 in PBS (PBST) for 30 minutes at room temperature, followed by overnight incubation with primary antibodies at the recommended dilution at 4°C. Primary antibodies used in this study included goat anti-Otx2 antibody (1:200, AF 1979; R&D systems), mouse anti-HuC/D (1:200, A21271; Thermo Fisher Scientific), rabbit anti-Sox9 antibody (1:500, AB5535; Millipore), rabbit anti-mCAR (1:200, AB15282, Millipore), rabbit anti-Nrl (1:150, AF2945, R&D Systems), mouse anti-PKCα (1:200, sc-8393, Santa Cruz), rabbit anti-Crx (1:500, PA5-32182, Thermo Fisher Scientific), goat anti-Sox2 (1:500, AF2018, R&D Systems), rabbit anti-Pax6 (1:500, AB2237, Millipore), rabbit anti-ZO1 (1:500, 61-7300, Thermo Fisher Scientific), rabbit anti-Rbpms (1:500, ab152101, Abcam). Following primary antibody incubation, the samples were washed thrice with PBST before incubation with a mixture of DAPI (0.5μg/ml) and secondary antibodies in the dark for 2 hours at room temperature.

Subsequently, the retinal sections were washed and mounted with an anti-fade solution before microscopy or storage. Slide images were captured using sing Nikon A1HD25 High speed and Large Field of View Confocal Microscope. Histological measurements and image processing were conducted using ImageJ software. The percentage of MG expressing various marker genes was quantified within retinal regions exhibiting near-complete AAV transduction.

### EdU Incorporation and Detection

5’-ethynyl-2’-deoxyuridine (EdU, 50mg/kg, Abcam ab146186) was intraperitoneally injected for 21 days after CCA administration to label the cells in the S phase. EdU staining was performed following the instruction of the Click-iT™ EdU Alexa Fluor™ 488 or 647 Imaging Kit (C10337, Thermo Fisher Scientific).

### *In situ* RNA hybridization

In the study, *in situ* RNA hybridization was conducted utilizing the RNAscope Multiplex Fluorescent Detection Reagents V2 kit (Advanced Cell Diagnostics) following standard commercial procedures. Initially, retinas were carefully dissected, fixed in 4% PFA, dehydrated using a sucrose solution, and embedded in OCT medium. Subsequently, the retinas were sectioned into 20μm slices and placed on SuperFrost Plus glass slides (Epredia).

After removing the OCT with PBS and dehydrating further with 50%, 70% and 100% ethanol, the retinal sections were incubated with a GFP antibody (AB_2307313; Aves Labs) overnight at 4°C. Post a triple wash with PBST (PBS with 0.1% Tween-20), the sections were hybridized with RNA probes: *Hes1* probe (Cat No.417701-C2 RNAscope™ Probe-Mm-*Hes1*-C2), *Pcdh17* probe (Cat No.489901-C2 RNAscope™ Probe-Mm-*Pcdh17*-C2) and Grm6 probe (Cat No.511611 RNAscope™ Probe-Mm-*Grm6*) for 2 hours at 40°C. Following the RNA hybridization process, the slides were stained with secondary antibodies (Jackson ImmunoResearch) and DAPI for two hours at room temperature.

The resulting fluorescent signals were observed and captured using a Nikon A1HD25 High-Speed and Large Field of View Confocal Microscope. The mRNA levels within the GFP-labeled nuclei membrane were then quantified by measuring the signal intensity level using ImageJ.

### MG sorting and snRNA library preparation and sequencing

Three or four fresh retinas were dissected from adult mice (strain: *Glast-Cre^ERT^;Rbpj^flox/flox^;Sun1:GFP* and *Glast-Cre^ERT^;Sun1:GFP*, aged as specified in Results) under RNase-free conditions in cold PBS on ice to minimize RNA degradation. Dissection tools (forceps, scissors) were sterilized with 70% ethanol and RNase Zap (Product #AM9780, Thermo Fisher Scientific) to eliminate RNase contamination. Retinas were frozen in liquid nitrogen until use. Retinal tissues were homogenized in 500µL of ice-cold NP-40 lysis buffer (Product #74385, Sigma-Aldrich) supplemented with 1× protease inhibitor cocktail (Product #P8340, Sigma-Aldrich) and 1U/µL RNase inhibitor (Product #EO0381, Thermo Fisher Scientific) using a pellet pestle (AST-YMB-15, Axyste). Homogenization was performed on ice with 15–20 gentle strokes to avoid excessive shearing of nuclei, followed by incubation on ice for 5 minutes to ensure complete lysis of cytoplasmic membranes while preserving nuclear integrity. The homogenate was filtered through a 70µm MACS® SmartStrainer (Catalog #130-110-915, Miltenyi Biotec) pre-rinsed with cold PBS to remove cell debris and intact cells. The filtrate (containing nuclei) was centrifuged at 500 × g for 5 minutes at 4°C to pellet nuclei, and the supernatant was discarded. The nuclear pellet was resuspended in 1mL of 1% BSA in PBS (1% BSA-PBS), supplemented with 1U/µL RNase inhibitor, to neutralize residual lysis buffer and stabilize nuclei. To enrich for MG nuclei (a rare population comprising 2–3% of total retinal cells), fluorescence-activated cell sorting (FACS) was performed using a Sony SH800Z cell sorter equipped with a 488-nm laser for GFP excitation and a 525/50-nm emission filter. Nuclei were gated based on forward scatter (FSC) and side scatter (SSC) to exclude debris and aggregates, and GFP-positive events were sorted into RNase-free 1.5mL tubes containing 100µL of 1% BSA-PBS with RNase inhibitor. Sorting was performed at 4°C with a flow rate of 1,000–2,000 events/second. Approximately 200,000 GFP+ nuclei were collected per sample and centrifuged at 500 × g for 5 minutes at 4°C. The pellet was resuspended in 1% BSA-PBS (with RNase inhibitor) and adjusted to a concentration of 600–800 nuclei/µL using a Countess II FL Automated Cell Counter (Thermo Fisher Scientific), ensuring compatibility with 10x Genomics Chromium systems for optimal Gel Bead-in-Emulsion (GEM) formation.

snRNA sequencing libraries were prepared using the Chromium Next GEM Single Cell 3’ GEM, Library & Gel Bead Kit v3.1 (Catalog #1000269, 10x Genomics), Chromium Next GEM Chip G Single Cell Kit (Catalog #1000127, 10x Genomics), and Chromium Controller iX. Briefly, 20µL of the nuclei suspension (containing ∼18,000 nuclei) was loaded into a well of the Chromium Next GEM Chip G, along with reverse transcription (RT) reagents, gel beads, and partitioning oil. GEMs were generated using the Chromium Controller iX, with each GEM containing a single nucleus, a gel bead (with barcoded oligonucleotides), and RT reagents. GEMs were transferred to 0.2-mL 8-tube strips (951010022, Eppendorf) and subjected to RT in a ProFlex™ PCR System (4484073, Thermo Fisher Scientific) with the following program: 53°C for 45 minutes, 85°C for 5 minutes, and hold at 4°C. Resulting cDNA was amplified using 11 cycles of PCR (98°C for 3 minutes; 12 cycles of 98°C for 15 seconds, 63°C for 20 seconds, 72°C for 1 minute; final extension at 72°C for 1 minutes) to generate sufficient material for library construction. Amplified cDNA was purified using SPRIselect beads (Beckman Coulter) at a 1:1.8 ratio (cDNA:beads) to remove primers and impurities. Purified cDNA was fragmented enzymatically (32°C for 5 minutes) using the Fragmentation Enzyme Mix (included in the 10x kit), followed by end repair, A-tailing, and ligation of Illumina-compatible adapters. Final libraries were purified with SPRIselect beads (1:1 ratio) and quantified using a Qubit 4 Fluorometer (Thermo Fisher Scientific).

Libraries were sequenced by Novogene (Beijing) on an Illumina NovaSeq 6000 platform using paired-end 150-bp reads. Each sample was allocated 100 Gb of sequencing data, ensuring an average depth of ∼35,000 reads per nucleus to capture robust transcriptomic profiles, including low-abundance transcripts relevant to MG function and differentiation. This expanded method provides detailed, reproducible parameters for each step, including reagent supplements, centrifugation conditions, FACS gating, and sequencing metrics, to support rigorous replication of the experiment.

### Preprocessing, filtering and clustering of snRNA data

The raw sequencing reads (paired-end 150 bp) were processed using Cell Ranger software (v6.1.2, 10x Genomics) following the manufacturer’s recommended pipeline (https://support.10xgenomics.com/single-cell-gene-expression/software/pipelines/latest/what-is-cell-ranger). For each sample, count was used to align reads to the mouse reference genome (mm10/GRCm38) and quantify gene expression. Cell Ranger’s built-in quality control metrics were used to generate filtered matrices, excluding potential cell-free RNA or debris-associated barcodes.

Subsequently, quality control, filtering, dimensional reduction, and clustering of the data were performed utilizing the Seurat package in R (*81*). Doublets and cells with less than 200 or more than 6000 expressed features and a percentage of mitochondrial transcripts exceeding 20% were excluded from further analysis. For the remaining cells, UMAP dimension reduction based on eight principal components (PCs) was implemented, and cells were clustered using the graphical clustering method within Seurat. Cell types were identified by leveraging known marker genes.

### snATAC library preparation and sequencing

Retinal MG nuclei were isolated as described in the snRNA-seq library preparation section. To permeabilize nuclear membranes for chromatin accessibility, isolated nuclei were treated with lysis buffer containing 5% digitonin (Thermo Fisher Scientific) for 1 minute on ice, then immediately washed and resuspended in diluted Nuclei Buffer (prepared by diluting 20X Nuclei Buffer, PN-2000207, 1:20 in nuclease-free water) to a final concentration of approximately 4,000 nuclei/µL.

snATAC-seq libraries were generated using the Chromium Next GEM Single Cell ATAC Kit v2 (PN-1000406, 10x Genomics) following the manufacturer’s protocol (CG000496 Rev C). In brief, permeabilized nuclei were incubated with Transposition Mix (ATAC Buffer B and ATAC Enzyme B) at 37°C for 30 minutes. Transposed nuclei were mixed with Master Mix (Barcoding Reagent B, Reducing Agent B, Barcoding Enzyme) and loaded onto a Chromium Chip H with ATAC Gel Beads v2 and Partitioning Oil to form GEMs using the Chromium Controller iX. GEMs were thermocycled (72°C 5 min; 98°C 30 sec; 12 cycles of 98°C 10 sec, 59°C 30 sec, 72°C 1 min; hold 15°C). Post-GEM cleanup used Dynabeads MyOne SILANE followed by double-sided SPRIselect purification. Indexed libraries were PCR-amplified for 7 cycles and purified again with SPRIselect. Library quality was assessed using an Agilent 2100 Bioanalyzer and sequenced by Novogene on an Illumina NovaSeq 6000 (PE150), with 120 Gb generated per sample.

### Preprocessing, Filtering and Clustering of snATAC-seq Data

The raw sequencing reads (paired-end 150 bp) were processed using Cell Ranger ATAC software (10x Genomics) following the manufacturer’s recommended pipeline. The workflow included: Illumina BCL files were converted to FASTQ format using cellranger-atac mkfastq, which assigned reads to individual samples based on index barcodes. For each sample, cellranger-atac count was used to align reads to the mouse reference genome (mm10/GRCm38) and generate fragment files quantifying chromatin accessibility.

Subsequently, quality control, filtering, dimensional reduction, and clustering of the data were performed utilizing the SnapATAC2 package in Python (*82*, *83*). Fragment files were imported and processed with sample-specific quality control parameters. Cells were filtered based on the following criteria: minimum fragment counts of 9,000, minimum transcription start site enrichment (TSS enrichment) score of 6, and maximum fragment counts of 90,000 to exclude low-quality nuclei and potential doublets. A tile matrix with 500 bp bins was generated, and the top 250,000 most variable features were selected for downstream analysis. Doublet detection was performed using the Scrublet algorithm implemented in SnapATAC2, and predicted doublets were removed from further analysis.

For dimensional reduction, spectral embedding (similar to Latent Semantic Indexing) was performed on the filtered cells, followed by UMAP for visualization. Cell clustering was performed using the Leiden algorithm after constructing a k-nearest neighbor graph. Cell types were identified by integrating gene activity scores derived from chromatin accessibility profiles with known marker genes and comparison to snRNA-seq data from matched samples.

### Electroretinography (ERG)

Mouse retinal function was assessed by ERG using the Espion E3 System (Diagnosys LLC), with a protocol adapted from previous studies on wild-type mice to characterize rod and cone responses (*10*). Mice were dark-adapted overnight, anesthetized with a ketamine/xylazine mixture (100/10 mg/kg), and their pupils were dilated with 5% phenylephrine (Mydrin-P, Santen) and 0.5% tropicamide for 5 min. Corneas were kept hydrated with gel. Gold-wire electrodes were placed on each cornea, with reference and ground electrodes in the mouth and tail, respectively. All procedures were performed under dim red light. Scotopic responses were elicited by 530 nm flashes of increasing intensity (0.01–30 cd·s/m²). For photopic recordings, mice were light-adapted for 5 min at 10 cd·s/m² to suppress rod activity, followed by flashes of 30 cd·s/m² on the same background. The amplitudes and implicit times of a- and b-waves were recorded for analysis.

### Optical Coherence Tomography (OCT)

In vivo OCT was performed using a Bioptigen SD-OCT system (Envisu R4310, Germany). Mice were anesthetized by intraperitoneal injection of ketamine/xylazine (100/10 mg/kg) and kept warm on a heating pad. Corneas and pupils were treated with 0.5% proxymetacaine (Provain-POS) and a mixture of 0.5% tropicamide and 0.5% phenylephrine (Mydrin-P, Santen), and hydrated with lubricating drops (Systane Ultra, Alcon) during imaging. Radial volume scans (1.7 mm diameter, 1000 A-scans/B-scan, 8 B-scans/volume, 24 frames/B-scan) were centered on the optic nerve. ONL thickness was measured 0.6 mm from the optic nerve head using ImageJ.

### Optomotor response test

Mouse visual acuity was assessed using an Optometry System (Cerebral Mechanics Inc.). Testing was performed with a grating drifting at 12 degrees/s and 100% contrast. The right eyes and left eyes were tested independently using counterclockwise and clockwise grating rotations, respectively. A staircase procedure was applied, with the observer progressing from low to high spatial frequencies to determine acuity thresholds. Each animal was tested for ∼10–15 min per session.

### Statistics

The data were expressed as mean ± standard error of the mean (s.e.m.). Sample sizes for each experiment were specified in the figure legend. Statistical analysis involved conducting one-way or two-way ANOVA followed by the Tukey test for comparing multiple groups, while the unpaired two-tailed Student’s t-test was employed for comparing two groups.

## Acknowledgements

This research was funded by Research Grants Council Hong Kong Project (11103819, 11102922, and 11100723), Hong Kong Health and Medical Research Fund Project (05160276 and 06172466), TUNG Biomedical Sciences Foundation, and Ming Wai Lau Center for Reparative Medicine Research Associate Program.

## Data availability

SnRNA-Sequencing data and snATAC-Sequencing data have been deposited in GEO under accession codes GSE331267.

## Author contributions

W.X. and B.L. conceived and designed the project. B.L. performed the experiments, collected and analyzed the data. B.L., Y.J., and S.L. carried out cell sorting and prepared the snRNA-seq and snATAC-seq libraries. L.C. and C.L. processed the Chromium Next GEM single-nucleus data using Cell Ranger. B.L. analyzed the snRNA-seq and snATAC-seq datasets, with assistance from C.L., B.L., W.W., H.T., and J.X. handled mouse work. B.L., J.Z. and Q.Z. optimized RNAscope imaging. B.L. wrote the original draft and organized the figures. W.X. reviewed and edited the manuscript and acquired funding.

---

**Figure S1.**
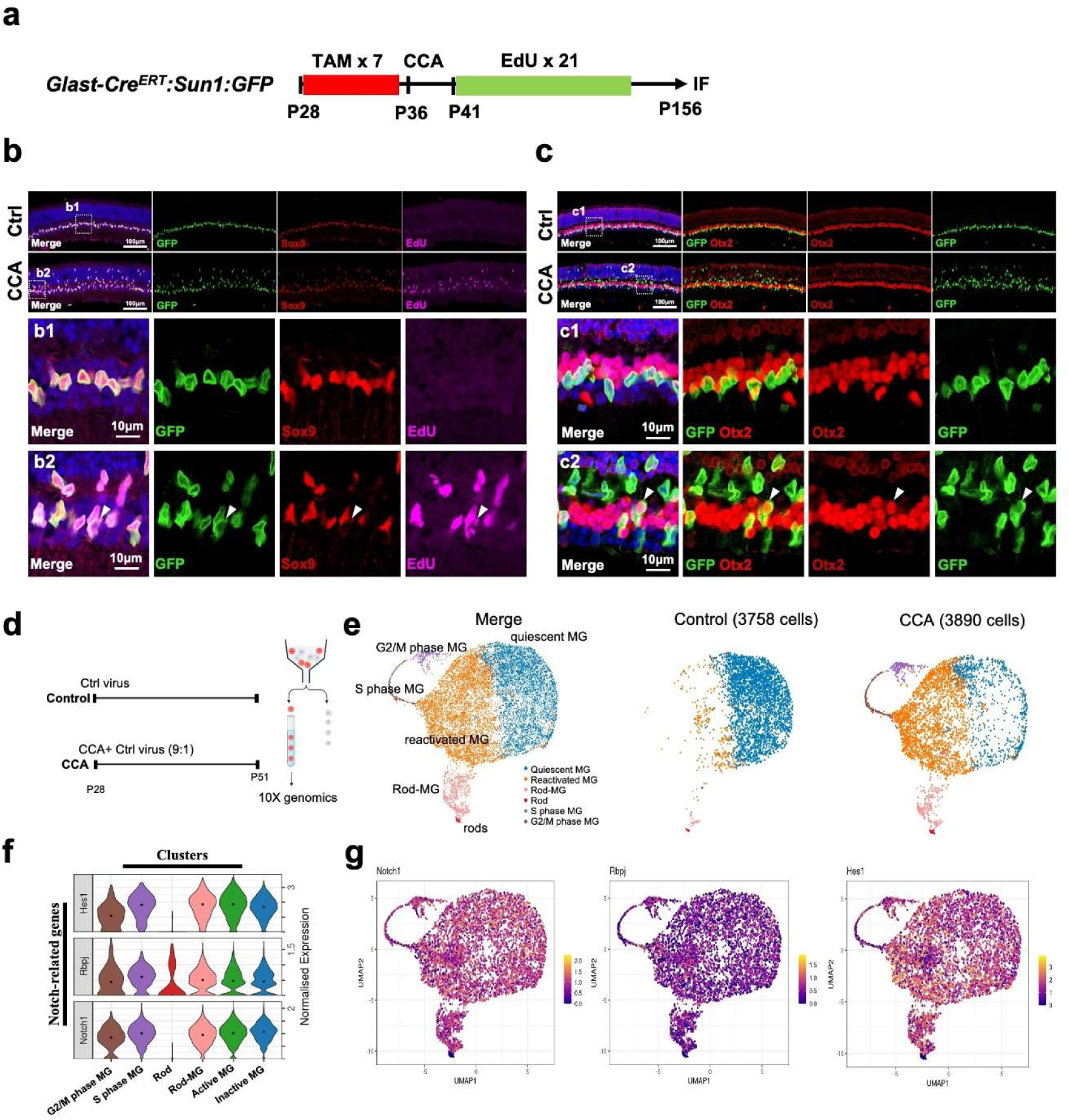
Notch signaling remains active in MG and MG-derived progeny. (a) Schematic illustrating the experimental design for labeling proliferating MG. (b) Representative Sox9 and EdU immunostaining on retinal sections from *Glast-Cre^ERT^;Sun1:GFP* mice treated with CCA at P28 and collected 4 months post-treatment. b1-b2 are the magnified views of the highlighted regions. The white arrows refer to GFP+ EdU+ Sox9- cells. (c) Representative Otx2 immunostaining on retinal sections from *Glast-Cre^ERT^;Sun1:GFP* mice treated with CCA at P28 and collected 4 months post-treatment. c1-c2 are the magnified views of the highlighted regions. The white arrows refer to GFP+ Otx2+ cells. (e) UMAP plot of scRNA-seq data for MG treated with CCA and control virus. (f) Split UMAP plots of the control and CCA groups. (g) Feature plots of normalized Notch related genes, *Notch1*, *Rbpj* and *Hes1* gene expression in different cell clusters.

**Figure S2.**
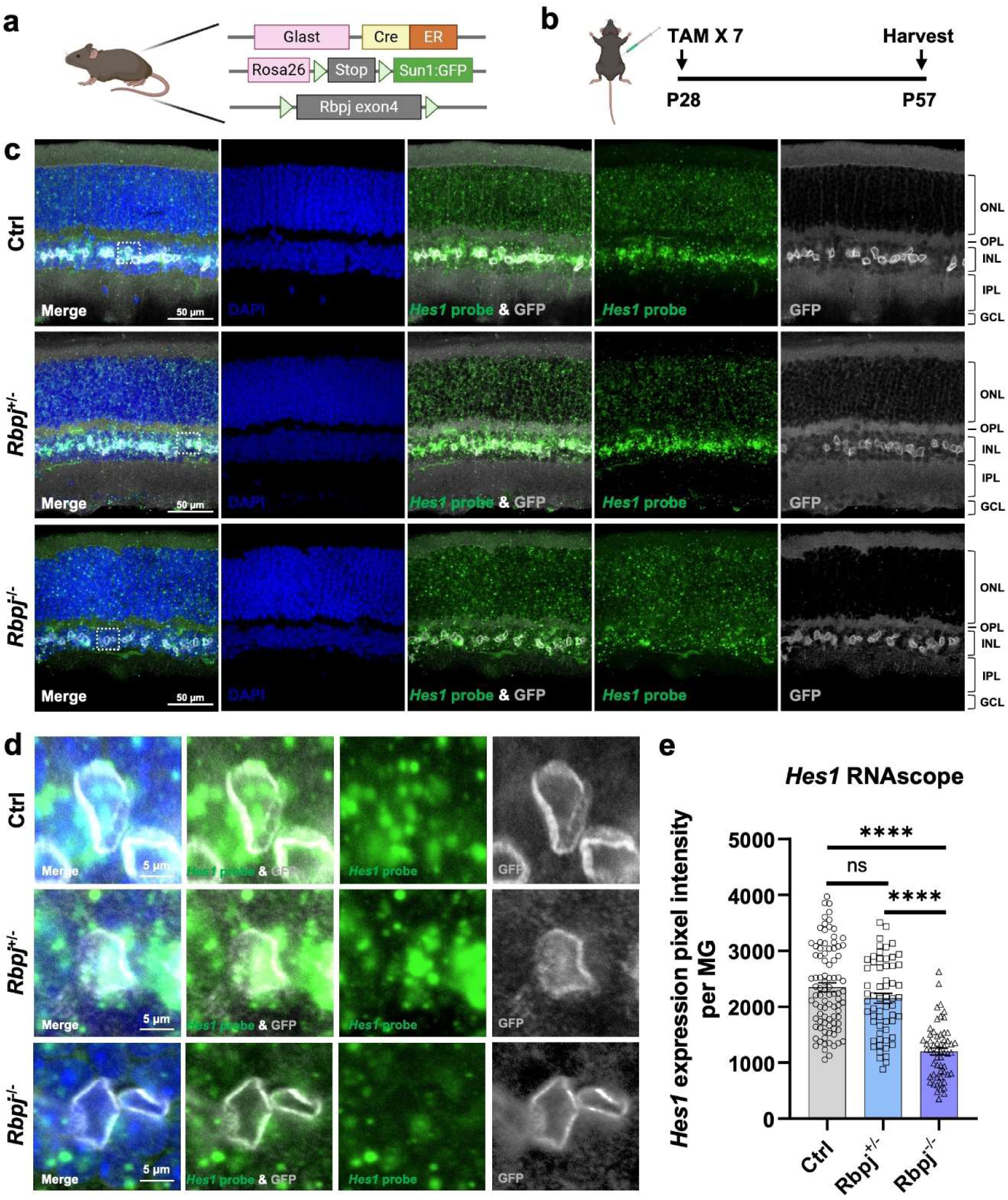
Deletion of *Rbpj* was sufficient to inhibit the canonical Notch signaling pathway. (a) Schematic of *Glast-Cre^ERT^;Rbpj^flox/flox^;tdT* mouse used in this study. (b) Schematic illustration of the examination of Notch inhibition via *Rbpj* deletion. (c) Hes1 mRNA in situ hybridization in *Glast-Cre^ERT^;tdT* (Ctrl), *Glast-Cre^ERT^;Rbpj^flox/wt^;tdT* and *Glast-Cre^ERT^;Rbpj^flox/flox^;tdT* mice received tamoxifen (TAM) injection. (d) Magnified views of the highlighted regions in (c). (e) The average pixel intensity of *Hes1* mRNA per GFP+ cell. n≥3 mice, data are presented as mean ± SEM. Ns=not significant, \*\*\*\**P* < 0.0001, by one-way ANOVA with Tukey’s post hoc test.

**Figure S3.**
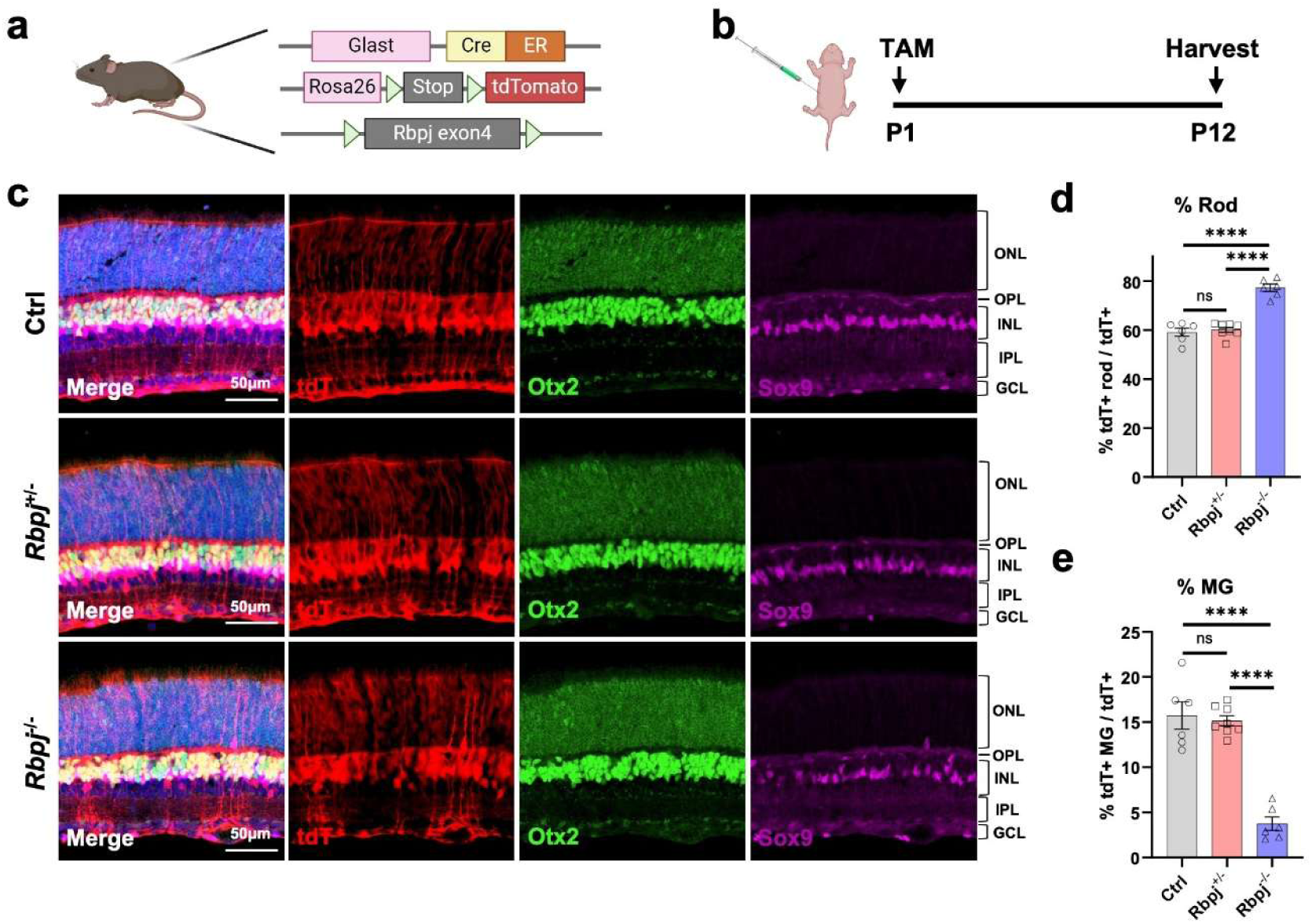
Overproduction of rod photoreceptor cells at expense of MG after Notch signaling inhibition. (a) Schematic of *Glast-Cre^ERT^;Rbpj^flox/flox^;Sun1:GFP* mouse used in this study. (b) Schematic illustration of the experiment examining how *Rbpj* deletion affects neurogenesis in retinal progenitor cells. *Glast-Cre^ERT^;tdT* (Ctrl), *Glast-Cre^ERT^;Rbpj^flox/wt^;tdT* and *Glast-Cre^ERT^;Rbpj^flox/flox^;tdT* mice were received TAM injection at postnatal day 1 (P1) and harvested at P12. (c) Representative immunostaining of Otx2 and Sox9 on retinal. (d) Percentage of tdT+ rod photoreceptor cells in overall tdT+ cells. (e) Percentage of tdT+ Sox9+ cells in overall tdT+ cells. n≥3 mice, ns. not significant, \*\*\*\**P* < 0.0001, by one-way ANOVA with Tukey post hoc test.

**Figure S4.**
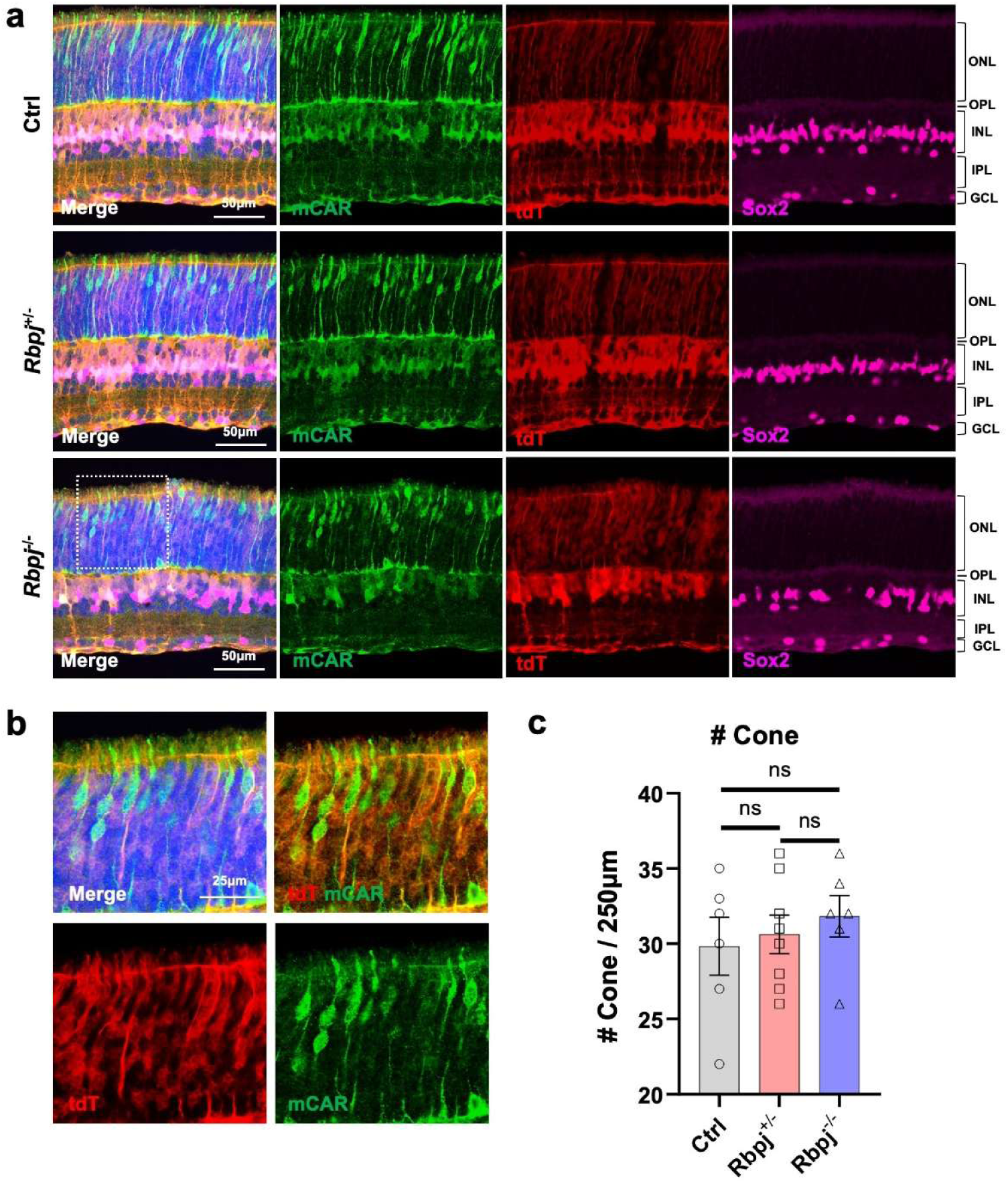
Deletion of *Rbpj* in late RPCs would not affect Cone photoreceptor cells formation. (a) Representative immunostaining of mCAR and Sox2 on retinal sections from *Glast-Cre^ERT^;tdT* (Ctrl), *Glast-Cre^ERT^;Rbpj^flox/wt^;tdT* and *Glast-Cre^ERT^;Rbpj^flox/flox^;tdT* mice received tamoxifen (TAM) injection at postnatal day 1 (P1) and harvested at P12. (b) Magnified views of the highlighted regions in (a). (c) Number of cone photoreceptor cells in 250μm. n≥3 mice, data are presented as mean ± SEM. ns=not significant, by one-way ANOVA with Tukey’s post hoc test.

**Figure S5.**
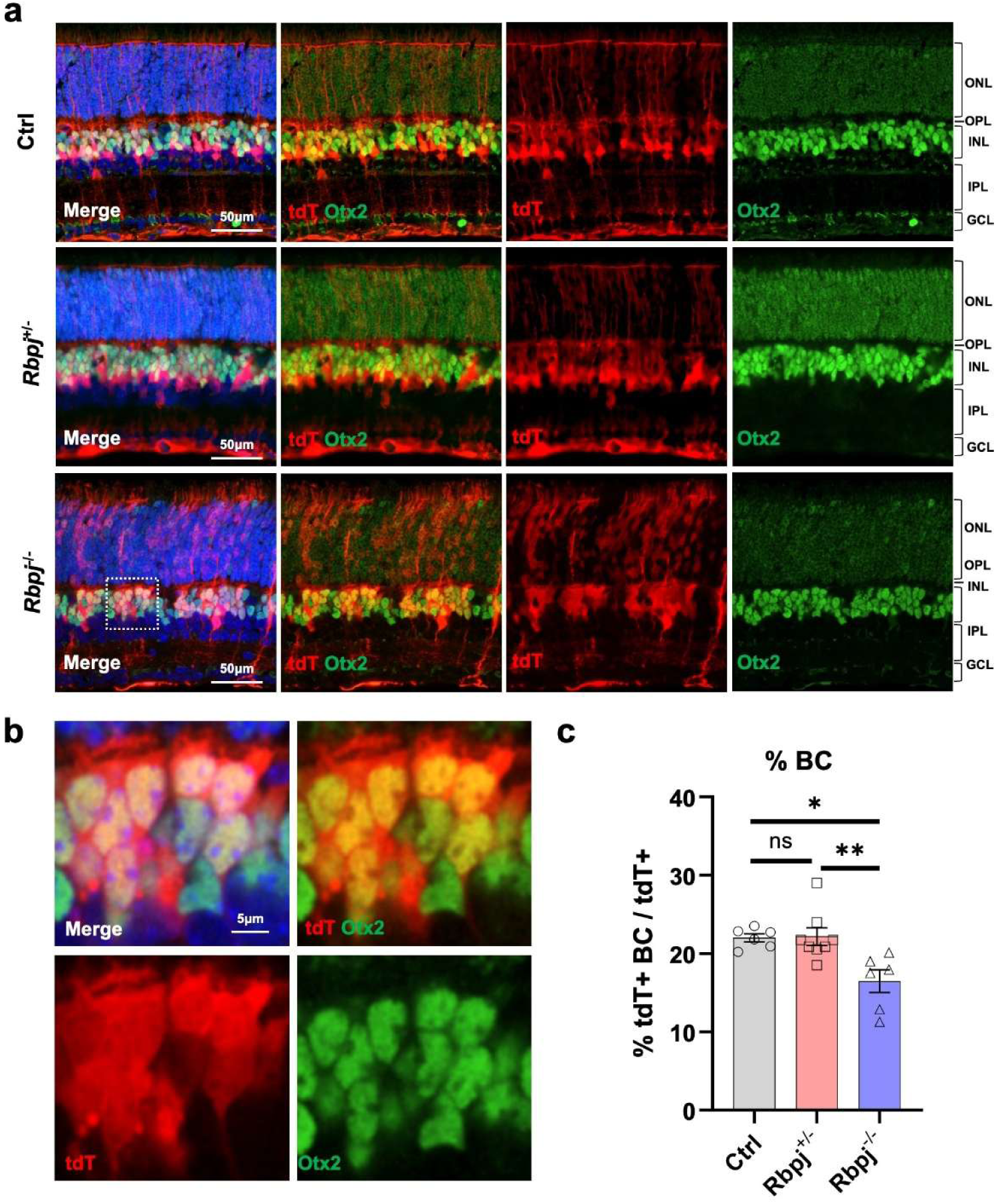
Notch signaling inhibition hindered the BCs generation from late RPCs. (a) Representative immunostaining of Otx2 on retinal sections from *Glast-Cre^ERT^;tdT* (Ctrl), *Glast-Cre^ERT^;Rbpj^flox/wt^;tdT* and *Glast-Cre^ERT^;Rbpj^flox/flox^;tdT* mice received tamoxifen (TAM) injection at postnatal day 1 (P1) and harvested at P12. (b) Magnified views of the highlighted regions in (a). (c) Percentage of tdT+ Otx2+ cells in overall tdT+ cells. n≥3 mice, ns=not significant, \**P* < 0.05, \*\**P* < 0.01, by one-way ANOVA with Tukey post hoc test.

**Figure S6.**
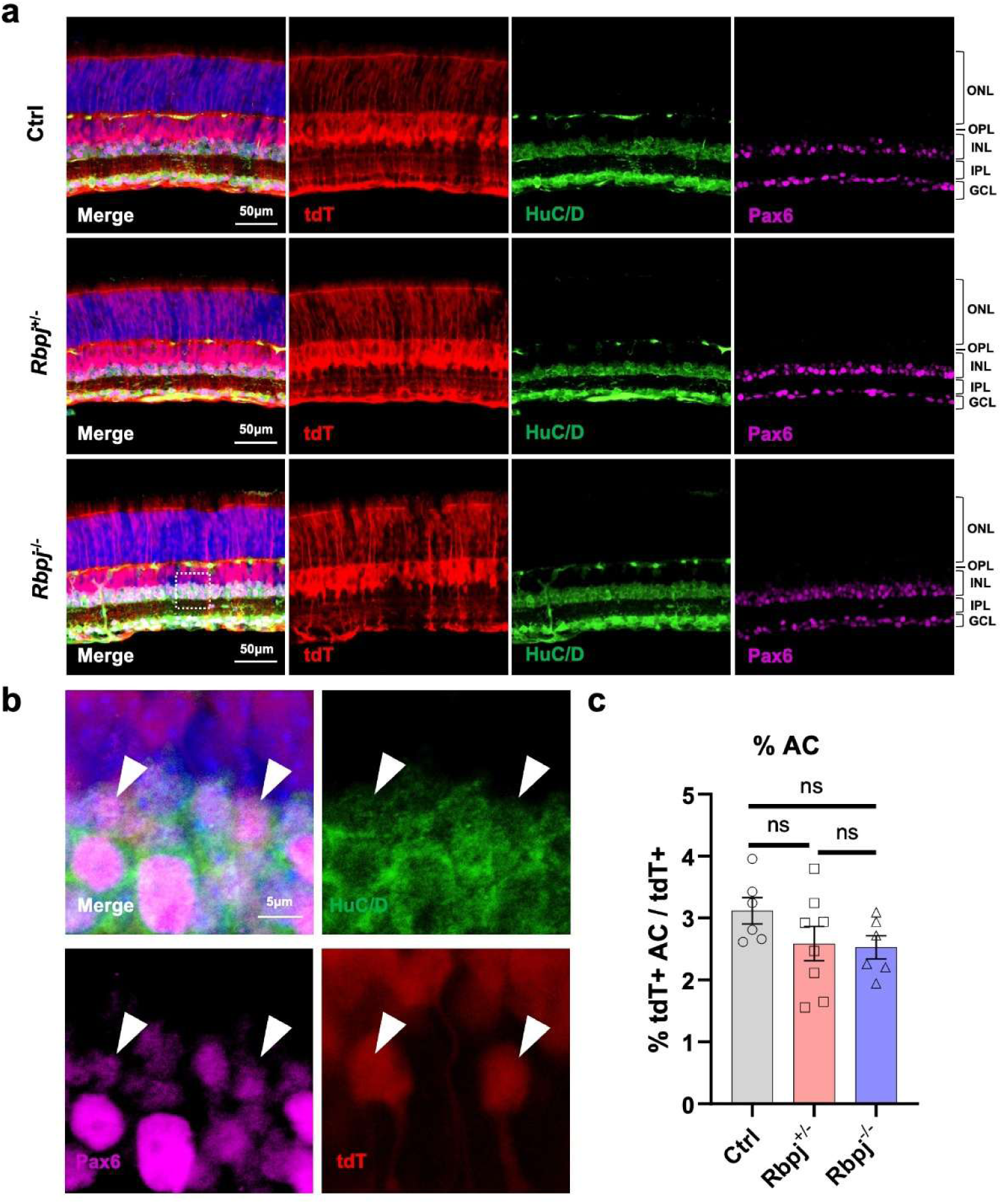
Deletion of *Rbpj* in late RPCs would not affect ACs formation. (a) Representative immunostaining of HuC/D and Pax6 on retinal sections from *Glast-Cre^ERT^;tdT* (Ctrl), *Glast-Cre^ERT^;Rbpj^flox/wt^;tdT* and *Glast-Cre^ERT^;Rbpj^flox/flox^;tdT* mice received TAM injection at postnatal day 1 (P1) and harvested at P12. (b) Magnified views of the highlighted regions in (a). (c) Percentage of tdT+ HuC/D+ Pax6+ cells in overall tdT+ cells. n≥3 mice, data are presented as mean ± SEM. ns=not significant, by one-way ANOVA with Tukey’s post hoc test.

**Figure S7.**
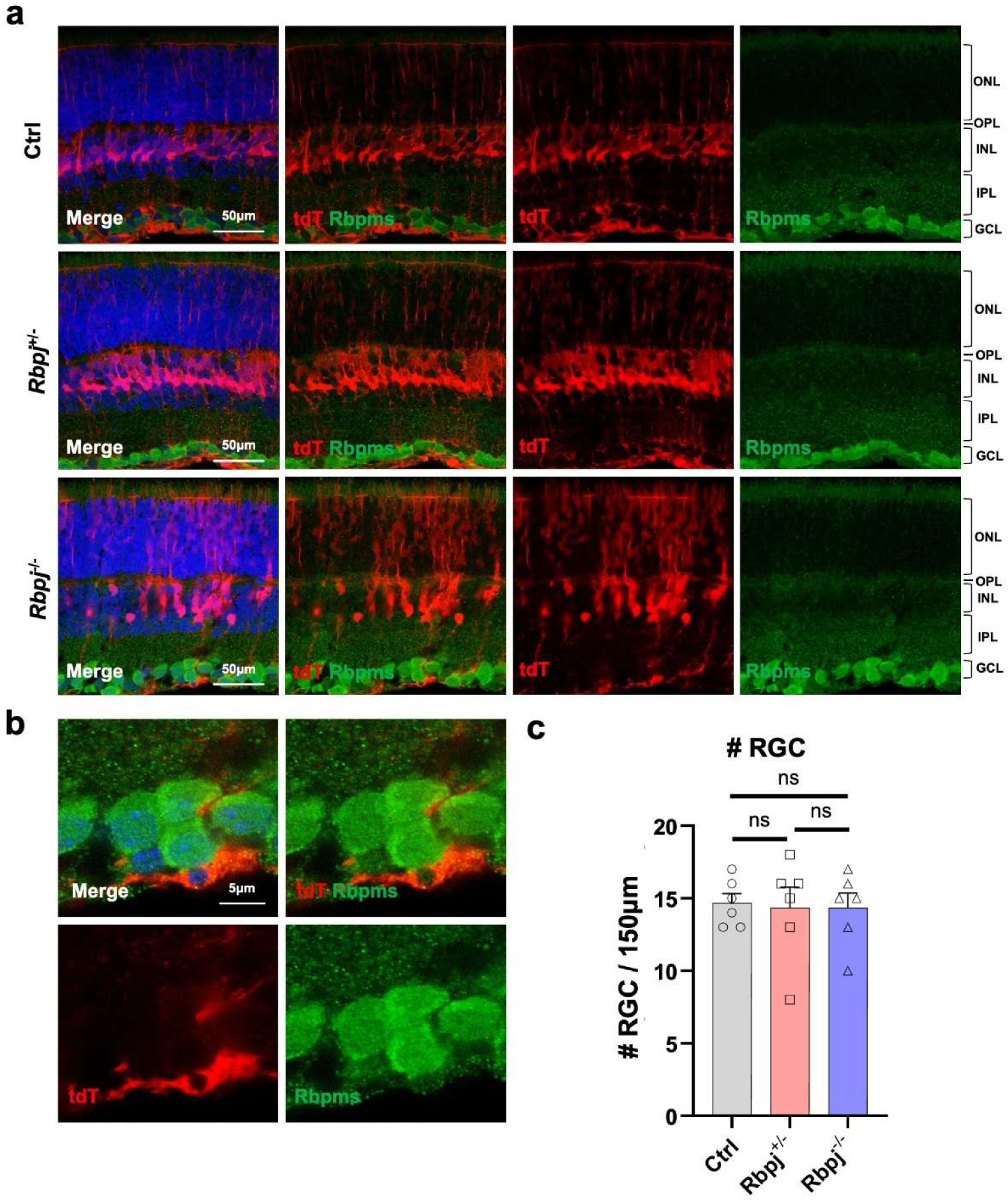
Deletion of *Rbpj* in late RPCs would not affect RGCs formation. (a) Representative immunostaining of Rbpms on retinal sections from *Glast-Cre^ERT^;tdT* (Ctrl), *Glast-Cre^ERT^;Rbpj^flox/wt^;tdT* and *Glast-Cre^ERT^;Rbpj^flox/flox^;tdT* mice received tamoxifen (TAM) injection at P1 and harvested at P12. (b) Magnified views of the highlighted regions in (a). (c) Number of Rbpms+ cells per 150μm. n≥3 mice, data are presented as mean ± SEM. ns=not significant, by one-way ANOVA with Tukey’s post hoc test.

**Figure S8.**
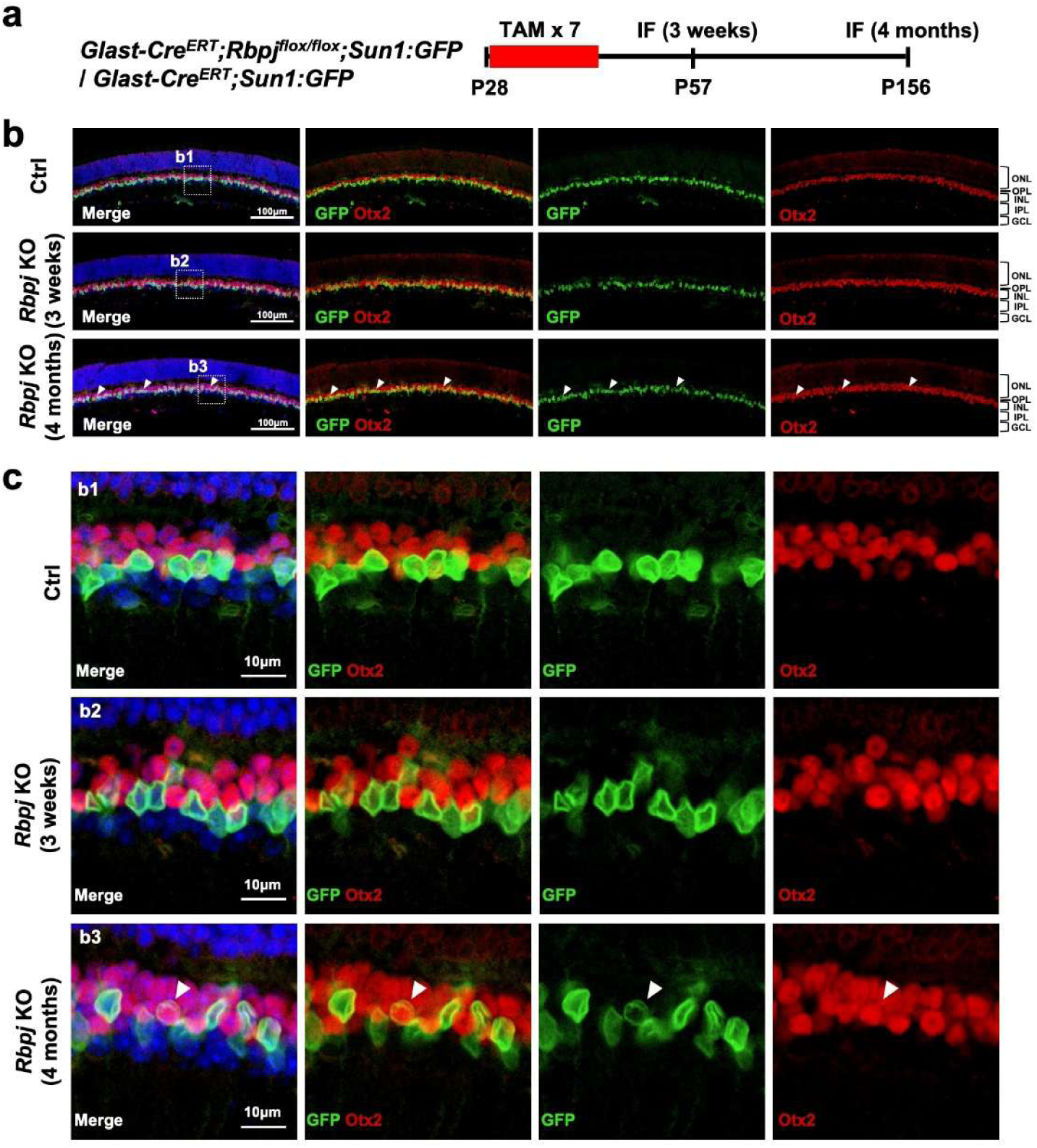
*Rbpj* deletion in adult MG induces limited neuronal conversion. (a) Schematic illustration of MG dedifferentiation and reprogramming experiment. (b) Representative immunostaining of Sox9 on retinal sections from *Glast-Cre^ERT^;Sun1:GFP* and *Glast-Cre^ERT^;Rbpj^flox/flox^;Sun1:GFP* mice harvested at different timepoints post TAM injection. The white arrows refer to GFP+ Sox9- cells. (c) Magnified views of the highlighted regions in (B).

**Figure S9.**
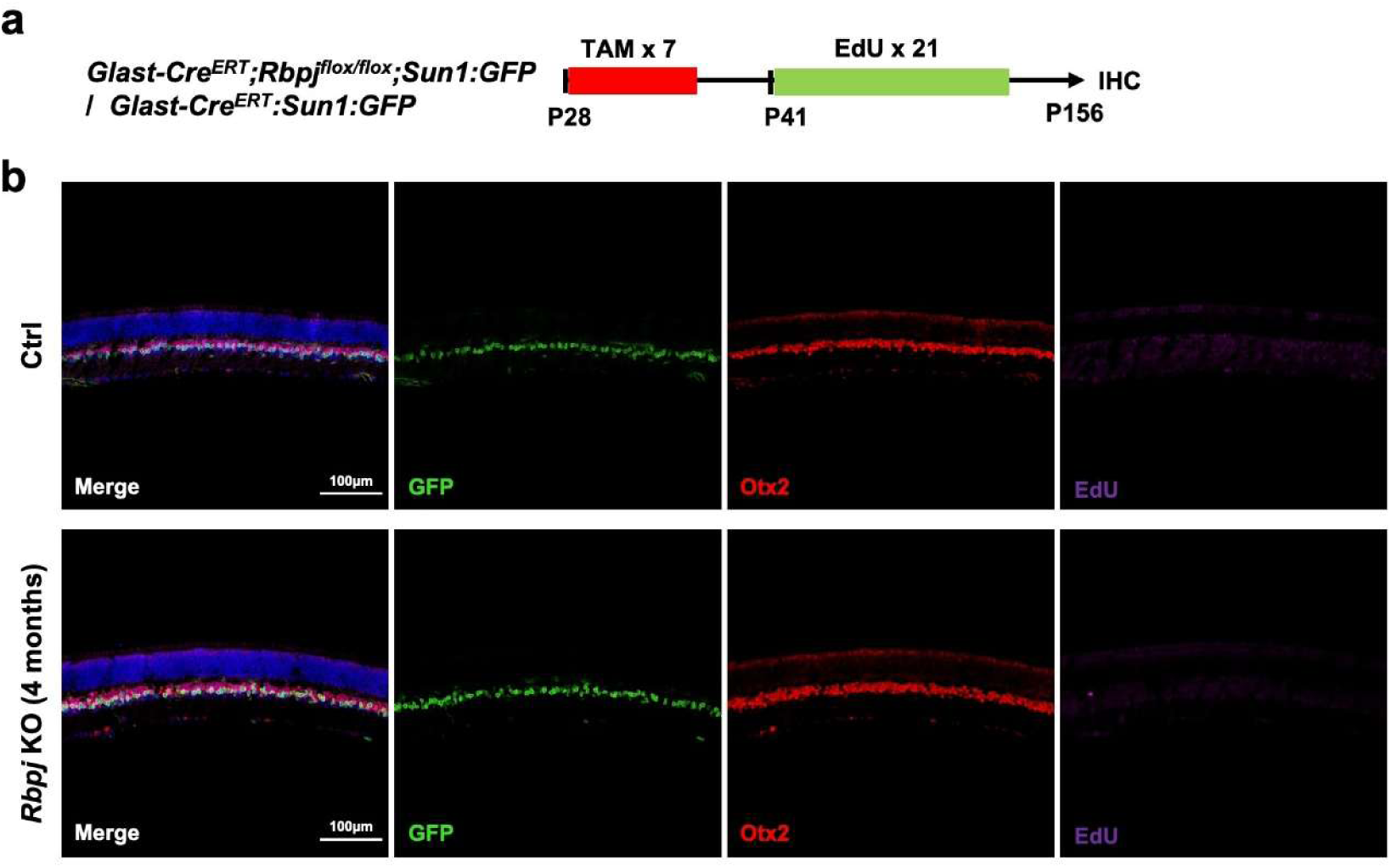
*Rbpj* deletion induces transdifferentiation-mediated neurogenesis in adult MG. (a) Schematic illustration of the experiment examining MG proliferation. (b) Representative immunostaining of Otx2 and EdU on retinal sections from *Glast-Cre^ERT^;Sun1:GFP* and *Glast-Cre^ERT^;Rbpj^flox/flox^;Sun1:GFP* mice harvested at 4 months post TAM injection.

**Figure S10.**
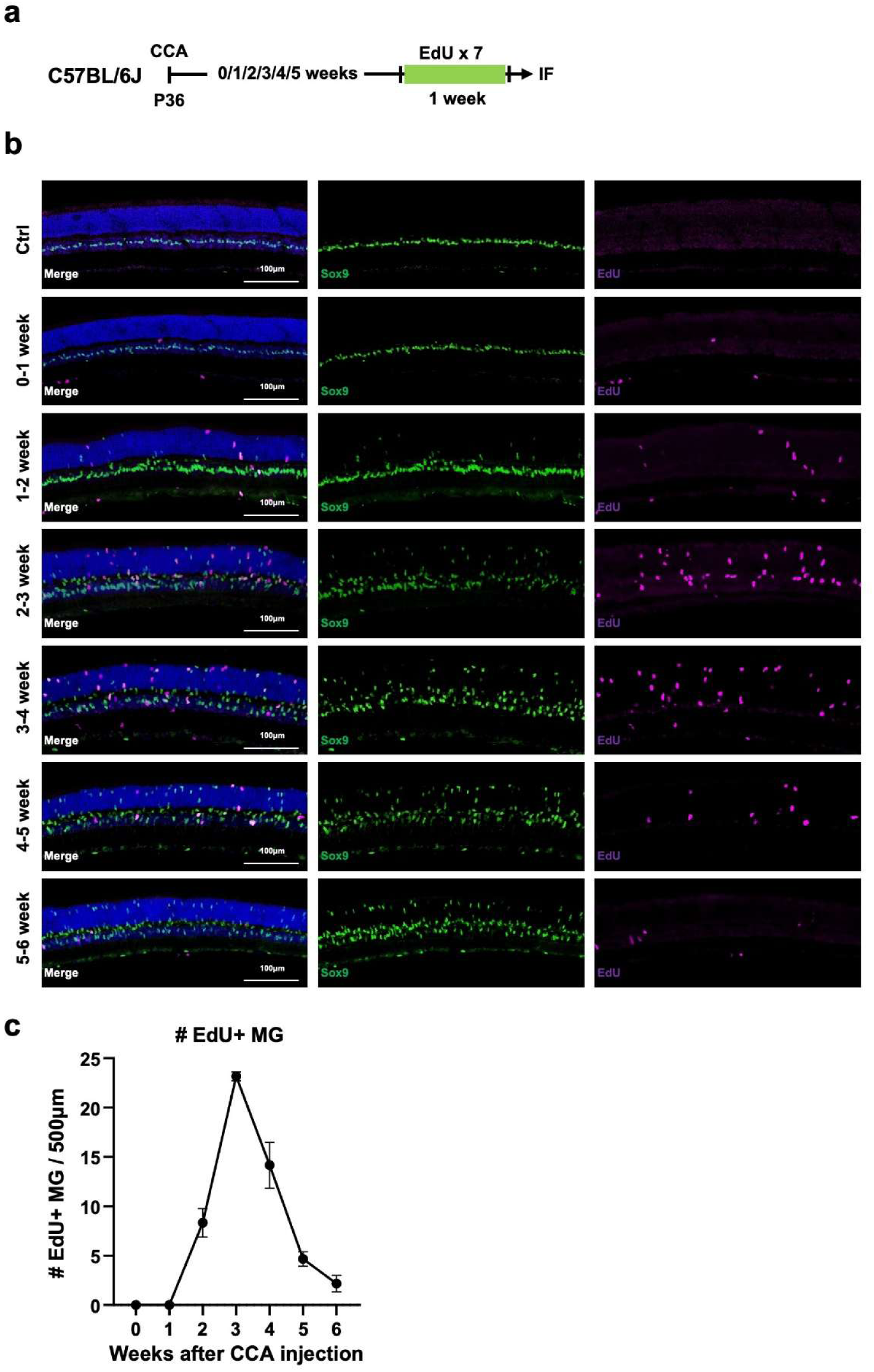
The self-limited process of MG proliferation induced by CCA. (a) Time-course analysis of MG proliferation following CCA injection. EdU was administered daily for 7 consecutive days starting at various weeks post-CCA injection, and retinas were harvested one day after the final EdU injection. (b) Representative EdU immunostaining from the time-course analysis of MG proliferation. (c) The changes in MG proliferation over time.

**Figure S11.**
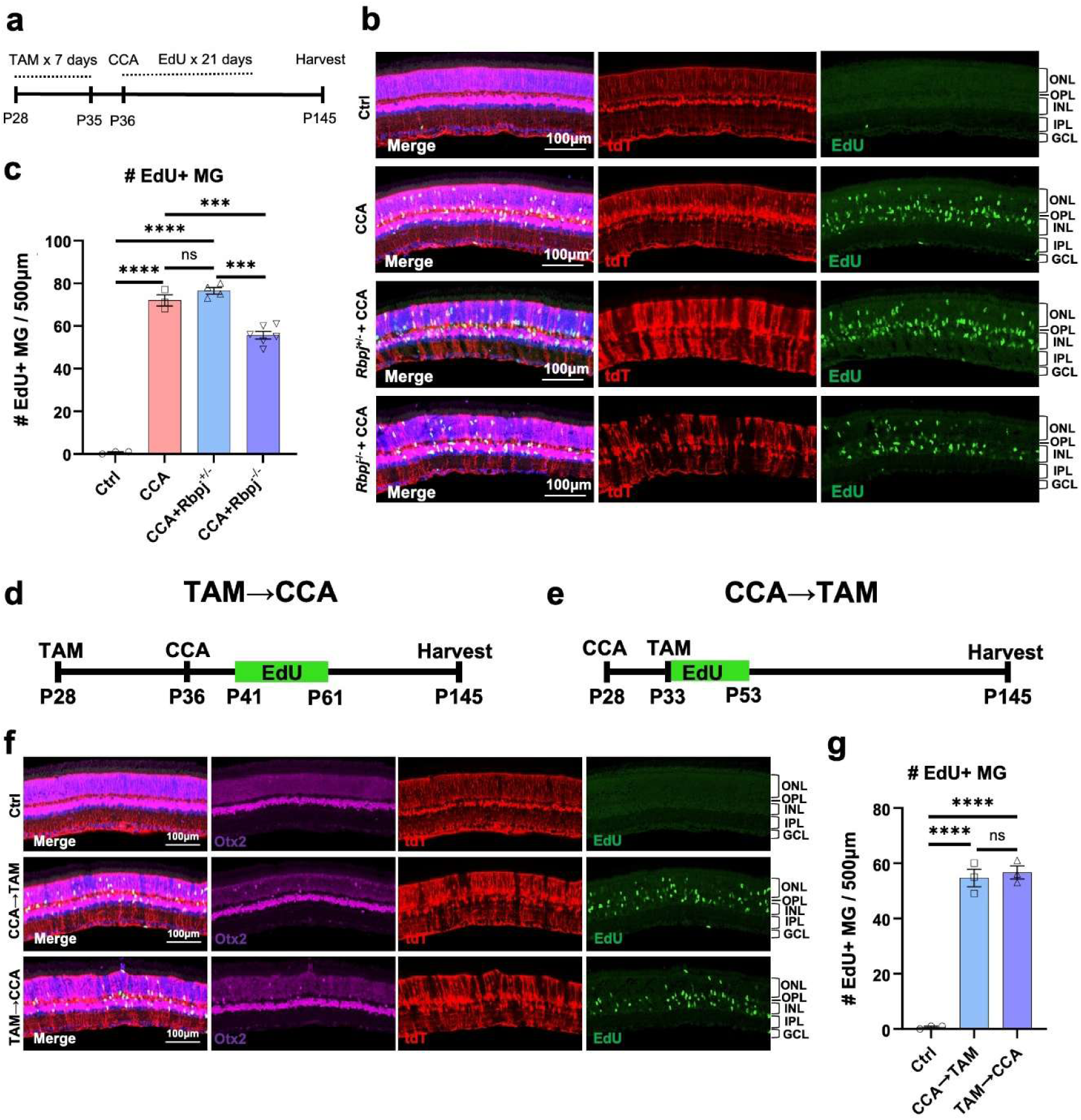
Notch inhibition reduced MG proliferation induced by CCA, yet it did not entirely prevent it. (a) Schematic illustration of proliferation level comparison experiment. (b) Representative immunostaining of EdU on retinal sections from *Glast-Cre^ERT^;tdT* (Ctrl), *Glast-Cre^ERT^;Rbpj^flox/wt^;tdT* and *Glast-Cre^ERT^;Rbpj^flox/flox^;tdT* mice received CCA injection. (c) Number of EdU+ cells in 500μm. n≥3 mice, data are presented as mean ± SEM. ns=not significant, \*\*\**P* < 0.001; \*\*\*\**P* < 0.0001, by one-way ANOVA with Tukey’s post hoc test. (d-e) Schematic illustration of the comparison of proliferation levels between different CCA injection and TAM administration orders. (f) Representative immunostaining of EdU on retinal sections from *Glast-Cre^ERT^;Rbpj^flox/flox^;tdT* mice received CCA injection. (g) Number of EdU+ cells in 500μm. N=3 mice, data are presented as mean ± SEM. Ns=not significant, \*\*\*\**P* < 0.0001, by one-way ANOVA with Tukey’s post hoc test.

**Figure S12.**
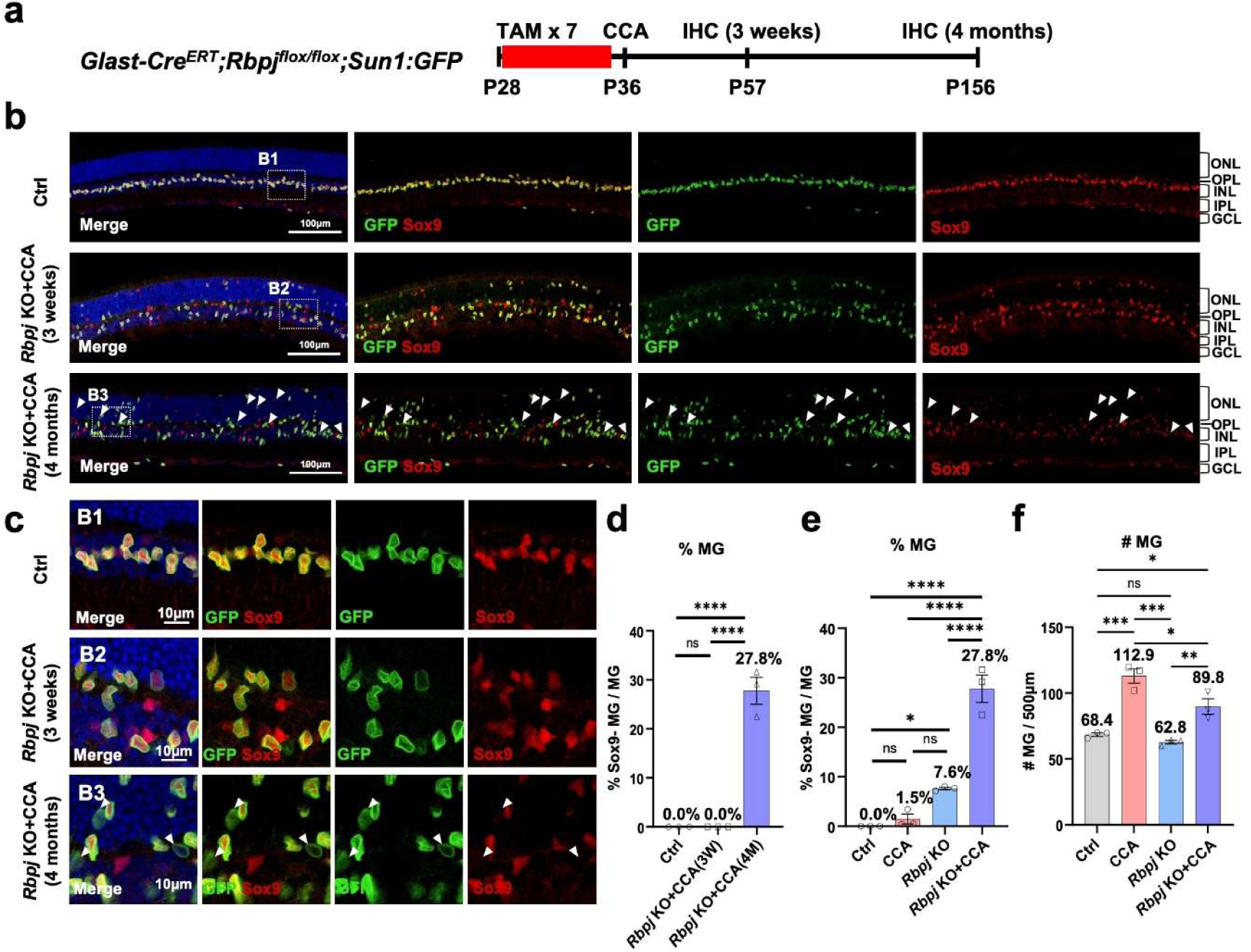
The synergistic effect of *Rbpj* KO and CCA resulted in a robust MG dedifferentiation. (a) Schematic illustration of MG dedifferentiation experiment. (b) Representative immunostaining of Sox9 on retinal sections from *Glast-Cre^ERT^;Sun1:GFP* (Ctrl) and *Glast-Cre^ERT^;Rbpj^flox/flox^;Sun1:GFP* mice at 3 weeks and 4 months post CCA treatment. The white arrows refer to GFP+ Sox9- cells. (c) Magnified views of the highlighted regions in (b). (d) Percentage of GFP+ Sox9- cells in overall GFP+ cells. 3W: 3 weeks, 4M: 4 months, n=3 mice, data are presented as mean ± SEM. ns=not significant, \*\*\*\**P* < 0.0001, by one-way ANOVA with Tukey’s post hoc test. (e) Percentage of GFP⁺ Sox9⁻ cells among total GFP⁺ cells in the 4-month samples. n=3 mice, data are presented as mean ± SEM. ns=not significant, \**P* < 0.05, \*\*\*\**P* < 0.0001, by one-way ANOVA with Tukey’s post hoc test. (f) Number of Sox9+ cells per 500μm in the 4-month samples. n=3 mice, data are presented as mean ± SEM. ns=not significant, \**P* < 0.05, \*\**P* < 0.01, \*\*\**P* < 0.001, by one-way ANOVA with Tukey’s post hoc test.

**Figure S13.**
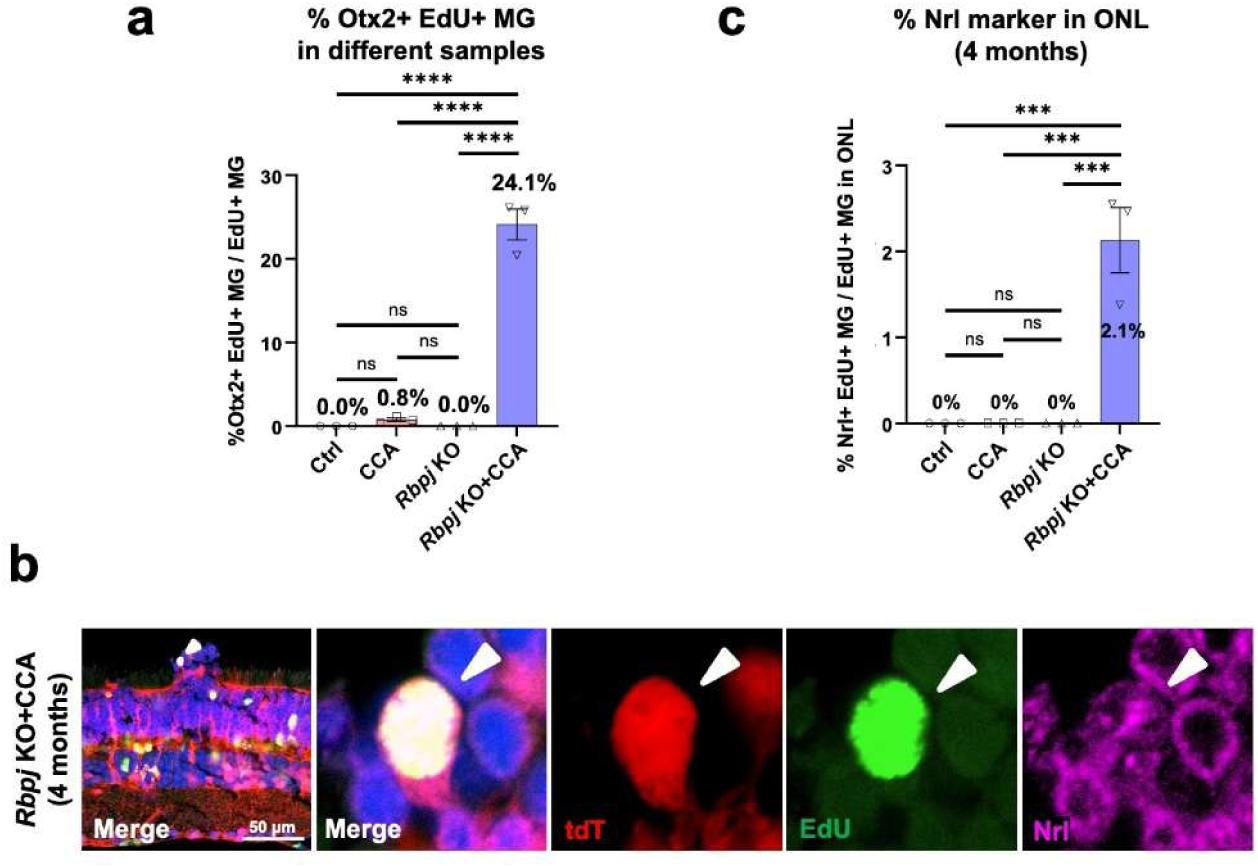
The synergistic effect of *Rbpj* KO and CCA resulted in a robust MG reprogramming. (a) Percentage of tdT+ EdU+ Otx2+ cells in overall tdT+ EdU+ cells. N=3 mice, data are presented as mean ± SEM. Ns=not significant, \*\*\*\**P* < 0.0001, by one-way ANOVA with Tukey’s post hoc test. (b) Representative immunostaining of EdU and Nrl on retinal sections. (c) Percentage of tdT+ EdU+ Nrl+ cells in ONL tdT+ EdU+ cells. N=3 mice, data are presented as mean ± SEM. Ns=not significant, \*\*\**P* < 0.001, by one-way ANOVA with Tukey’s post hoc test.

**Figure S14.**
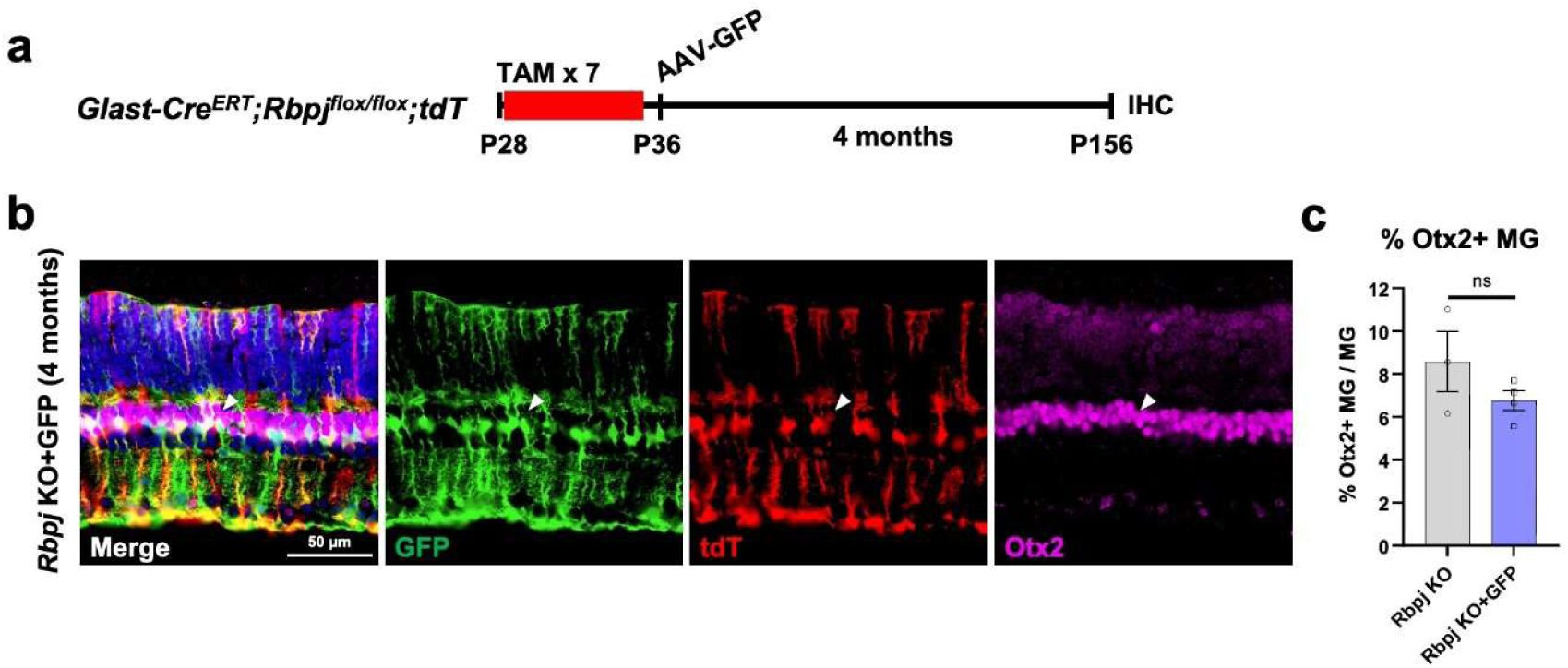
Control AAV does not enhance MG reprogramming efficiency. (a) Schematic of AAV7m8-GFAP-GFP-WPRE injection used as a control to exclude nonspecific effects of AAV administration. (b) Representative immunostaining of Otx2 on retinal sections. The white arrows refer to tdT+ Sox9+ GFP- cells. (c) Percentage of tdT+ Otx2+ cells in overall tdT+ cells. Each data point represents one mouse, data are presented as mean ± SEM. Ns=not significant, by unpaired two-tailed student’s t-test.

**Figure S15.**
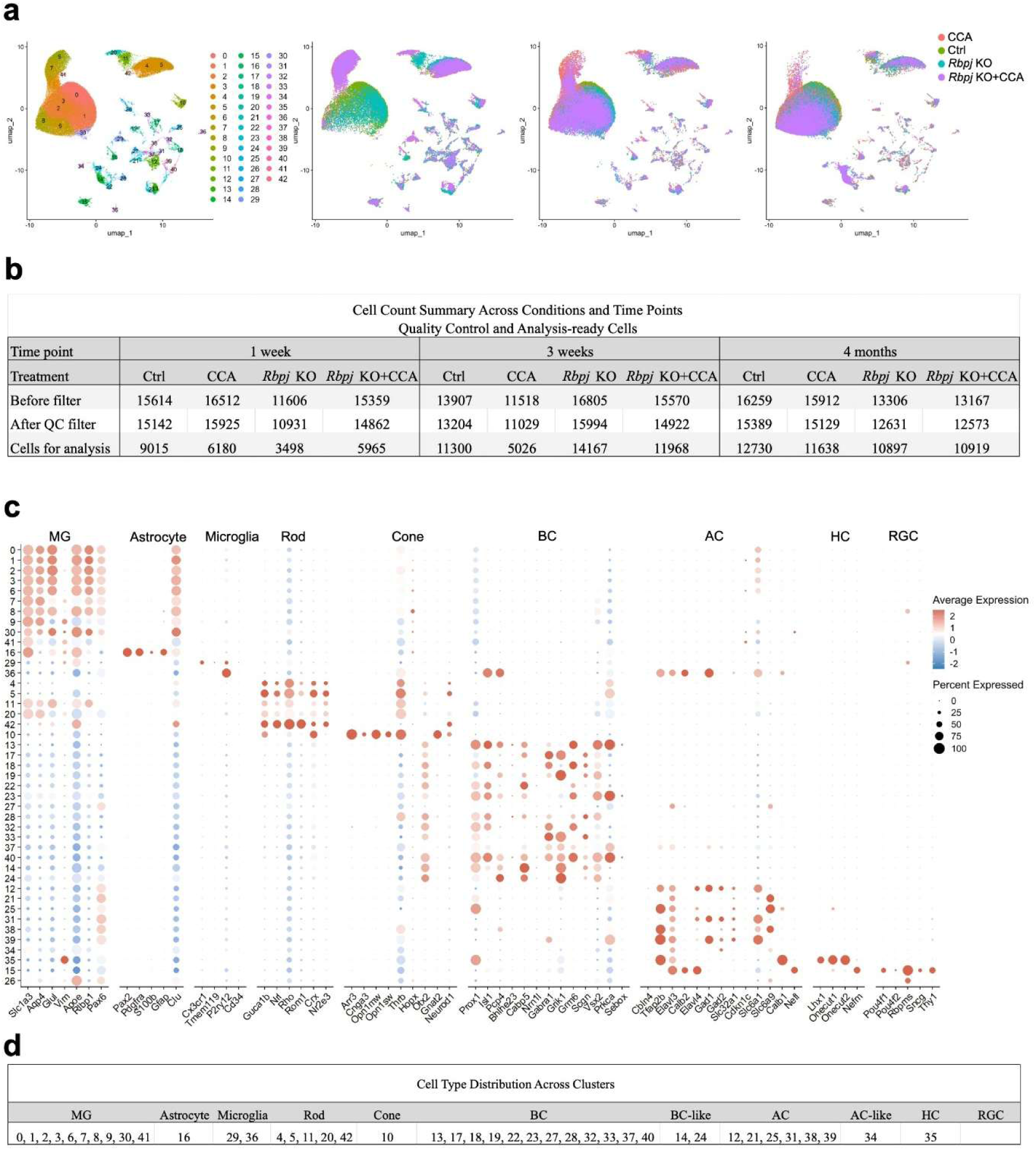
Preprocessing and filtering of snRNA data. (a) The UMAP before the removal of contamination cells. The original mature neurons affecting the analysis were removal. (b) Number of cells passing quality control and used for snRNA-seq analysis. (c) Dot plot showing the expression of marker genes of each cluster in unfiltered UMAP. (d) The initial annotation of clusters in the unfiltered UMAP.

**Figure S16.**
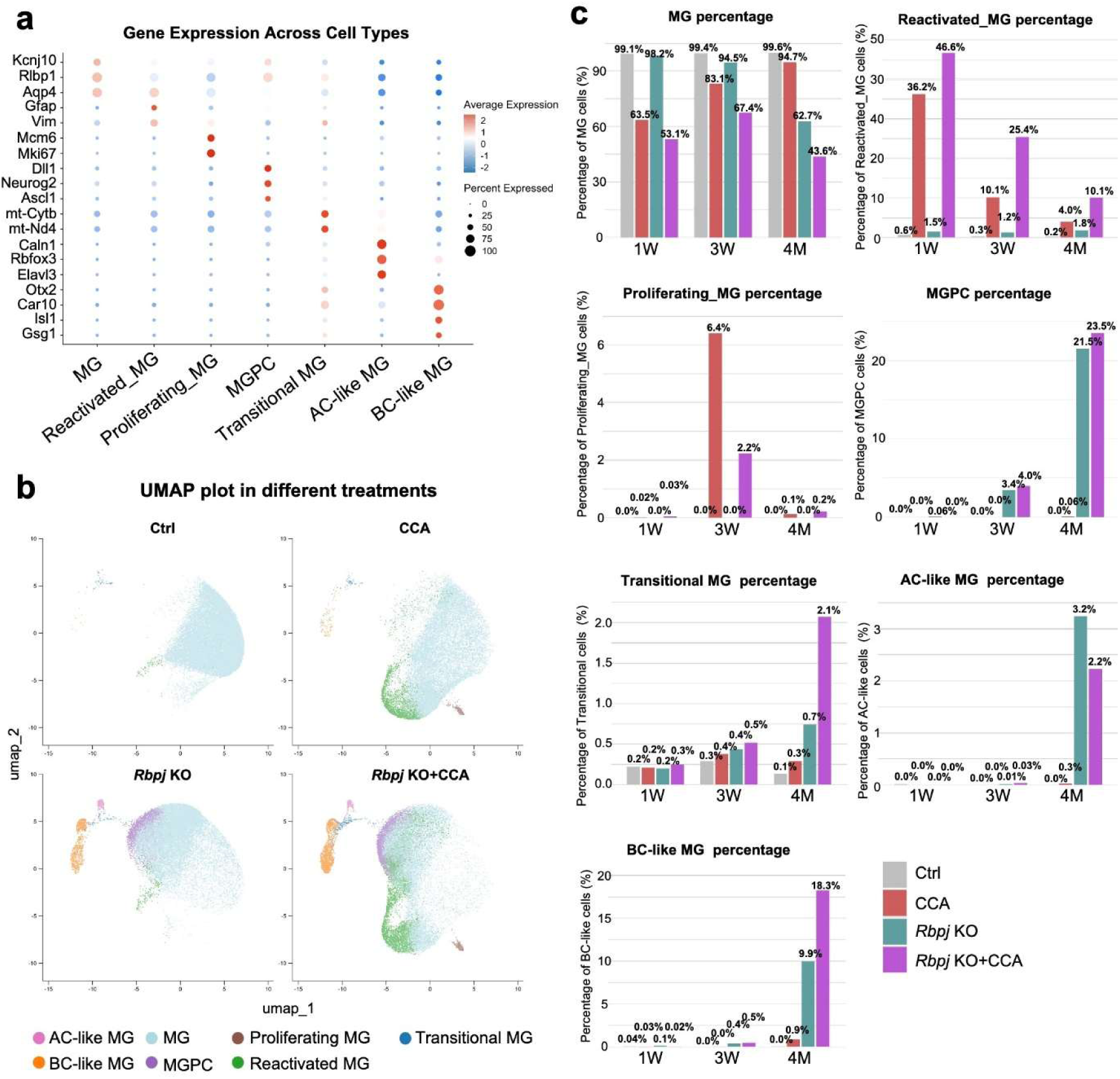
Additional snRNA-seq analysis of MG at 1 week, 3 weeks and 4 months post-CCA treatment. (a) Dot plot showing gene expression and cell percentages for quiescent MG, proliferating MG, reactivated MG, MGPC, transitional MG, AC-like MG, BC-like MG. (b) Separated UMAP plot of snRNA-seq data of Ctrl, CCA, *Rbpj* KO, *Rbpj* KO+CCA treatment. (c) Separated proportions of cell clusters within Ctrl, CCA, *Rbpj* KO and *Rbpj* KO+CCA groups at different timepoints.

**Figure S17.**
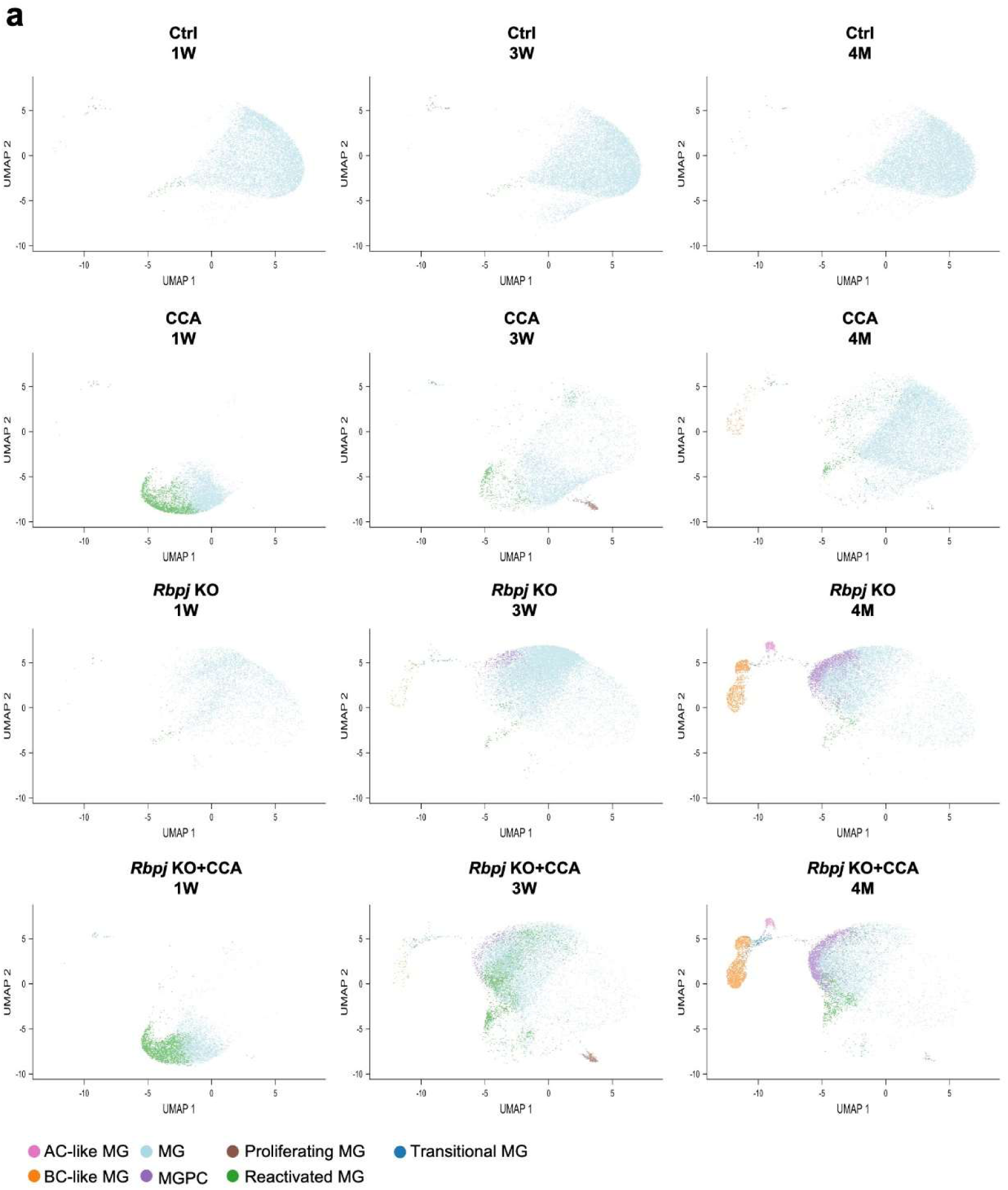
Separation of UMAP by different timepoints and treatments. (a) The split UMAP by the condition of time and treatment. To monitor the progress of MG regeneration, we implemented three time points (1 week (1W), 3 weeks (3W), 4 months (4M)) and four treatment groups (Ctrl, CCA, *Rbpj* KO, *Rbpj* KO+CCA).

**Figure S18.**
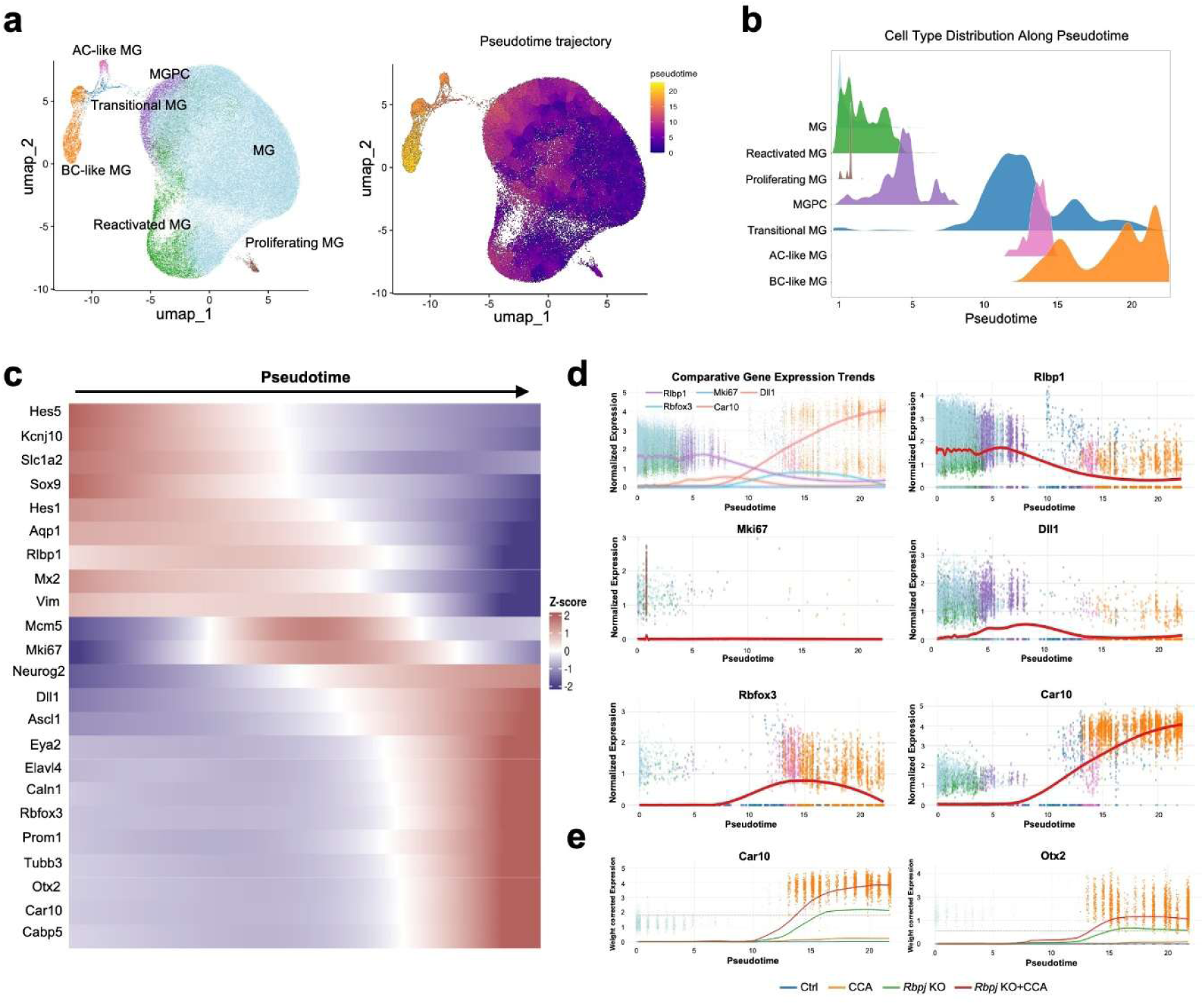
Pseudotime trajectory analysis of MG reprogramming process. (a) Pseudotime trajectory analysis showing the MG reprogramming process. (b) Dynamic cell type distribution along the pseudotime. (c) Change of gene expression level along the pseudotime. MG (Hes5, Kcnj10, Slc1a2, Sox9, Hes1, Aqp1, Rlbp1); Reactivated MG (Mx2, Vim); Proliferating MG (Mcm5, Mki67); MGPC (Neurog2, Dll1, Ascl1, Eya2); AC-like (Elavl4, Caln1, Rbfox3); photoreceptor cell (Prom1); RGC (Tubb3); BC-like (Otx2, Car10, Cabp5). (d) Gene expression trends along pseudotime. (e) Separated gene expression trends along pseudotime.

**Figure S19.**
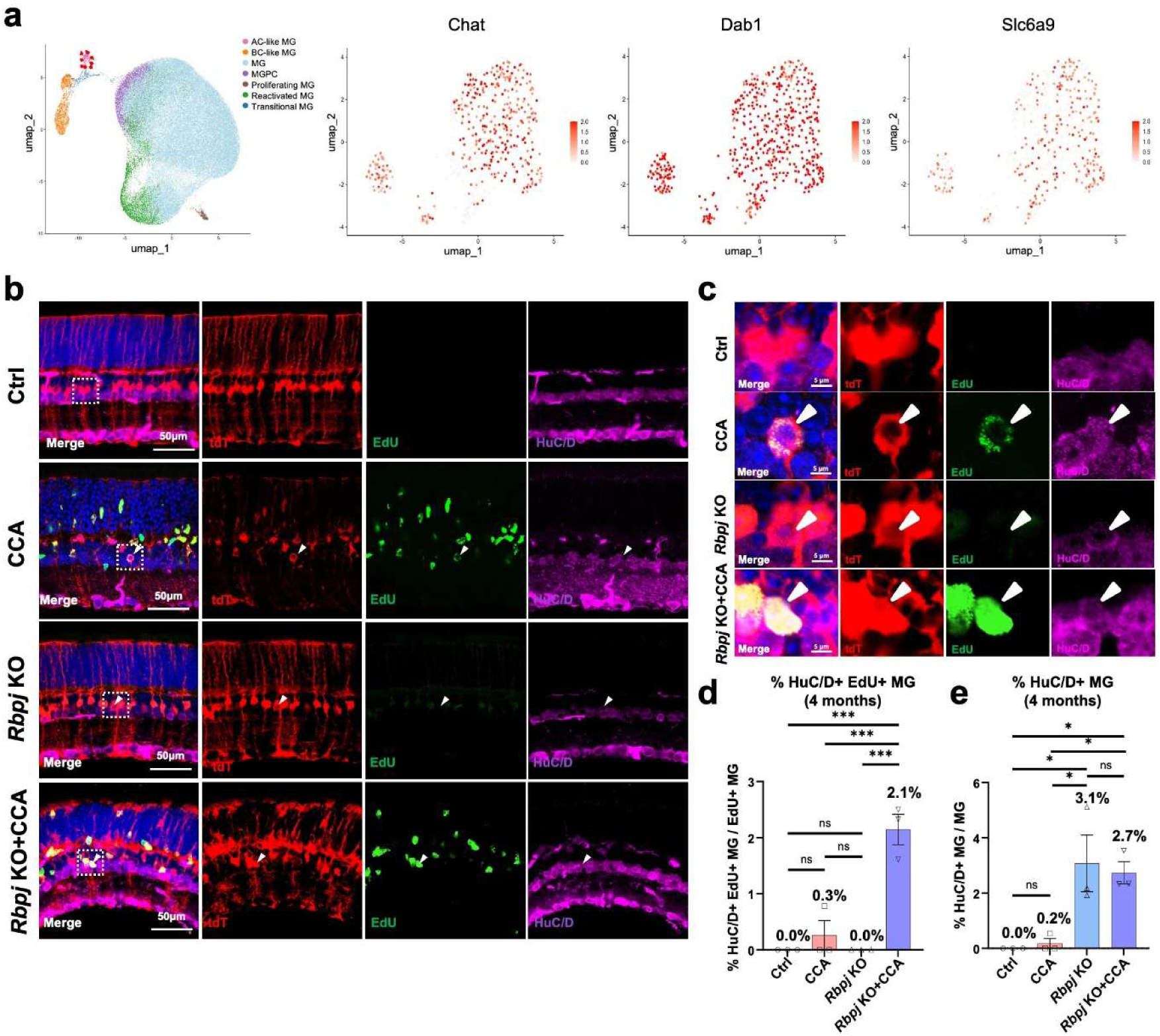
Subclustering analysis of the AC-like population. (a) Feature plots of AC-like subtypes showing the expression patterns of Chat (starburst ACs), Dab1 (A17 ACs), and Slc6a9 (nGnG ACs). The AC-like population is outlined in red. (b) Representative immunostaining of EdU and HuC/D on retinal sections. The white arrows refer to tdT+ HuC/D+ cells. (c) Magnified views of the highlighted regions in (b). (d) Percentage of tdT+ EdU+ HuC/D+ cells in overall tdT+ EdU+ cells. n=3 mice, data are presented as mean ± SEM. ns=not significant, ***P < 0.001, by one-way ANOVA with Tukey’s post hoc test. (e) Percentage of tdT+ HuC/D+ cells in overall tdT+ cells. n=3 mice, data are presented as mean ± SEM. Ns=not significant, *P < 0.05, by one-way ANOVA with Tukey’s post hoc test.

**Figure S20.**
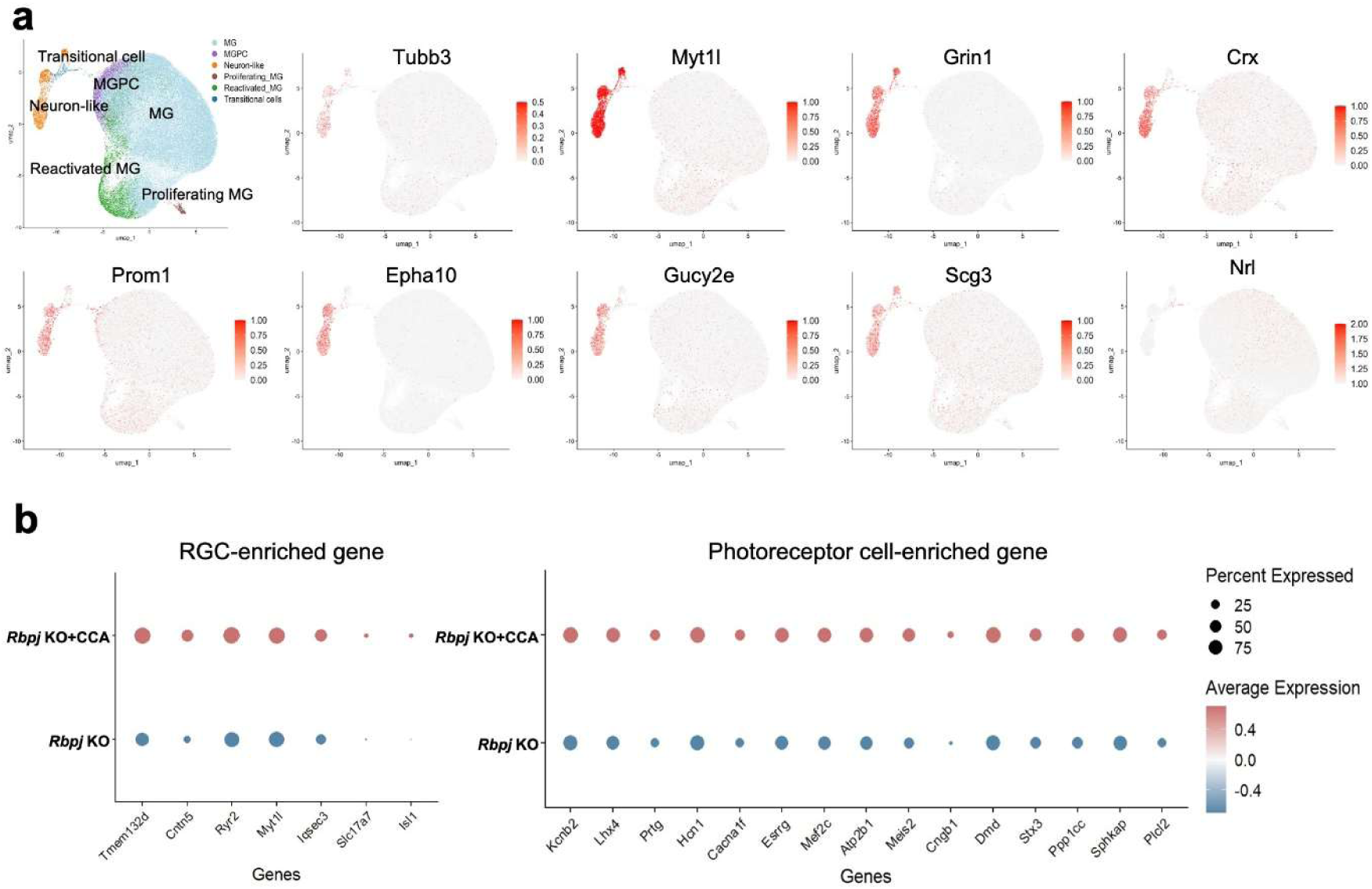
The newborn neurons showed diverse gene expression profiles linked to different neuronal cell types. (a) Feature plots showing the additional neuronal markers related to RGC and photoreceptor cells. (b) Dot plots showing different expression of RGC-enriched and photoreceptor cells-enriched genes (p<0.05).

**Figure S21.**
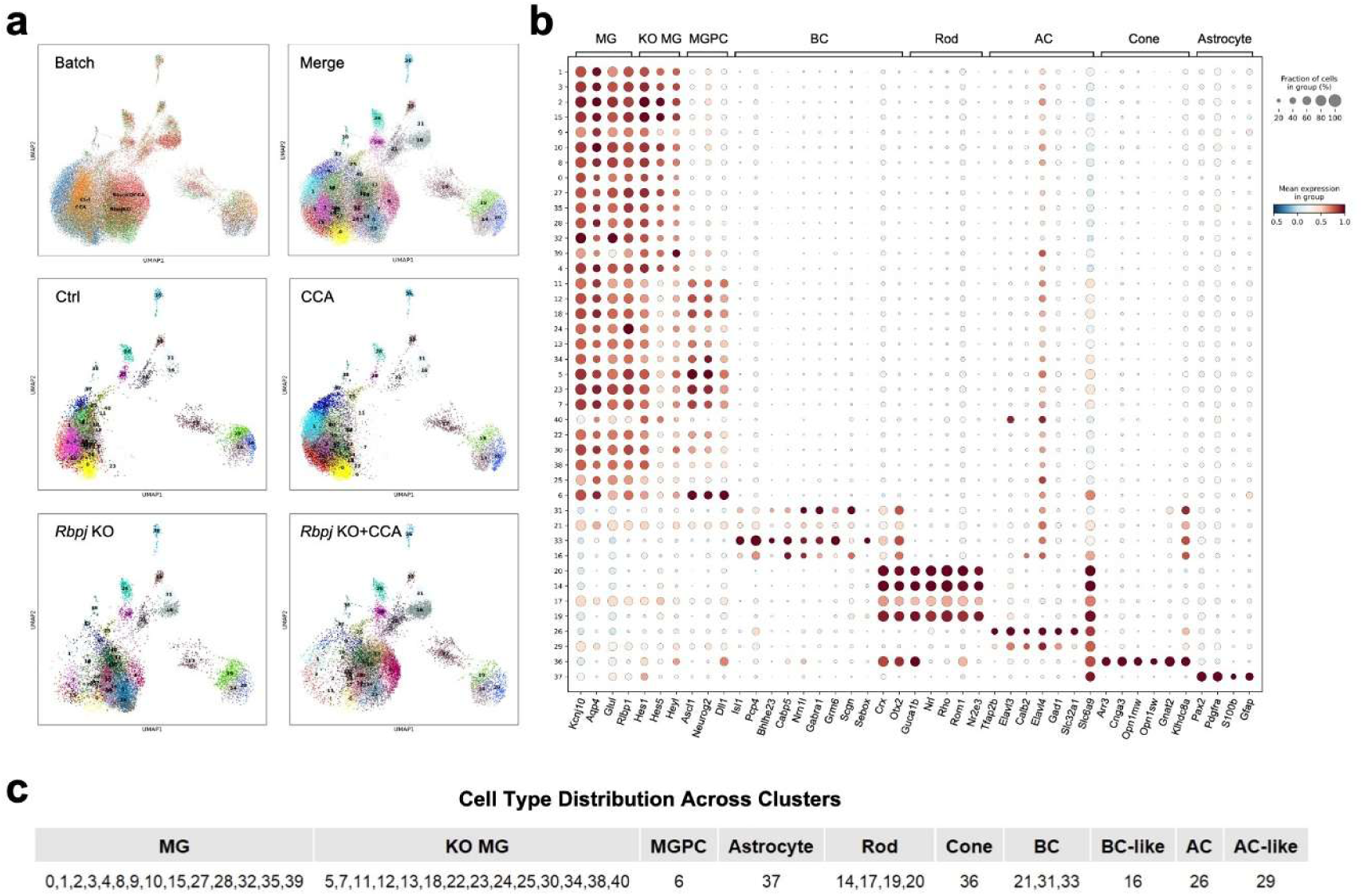
Preprocessing and filtering of snATAC data. (a) The UMAP before the removal of contamination cells. (b) Dot plot showing the expression of marker genes of each cluster in unfiltered UMAP. (c) The initial annotation of clusters in the unfiltered UMAP.

**Figure S22.**
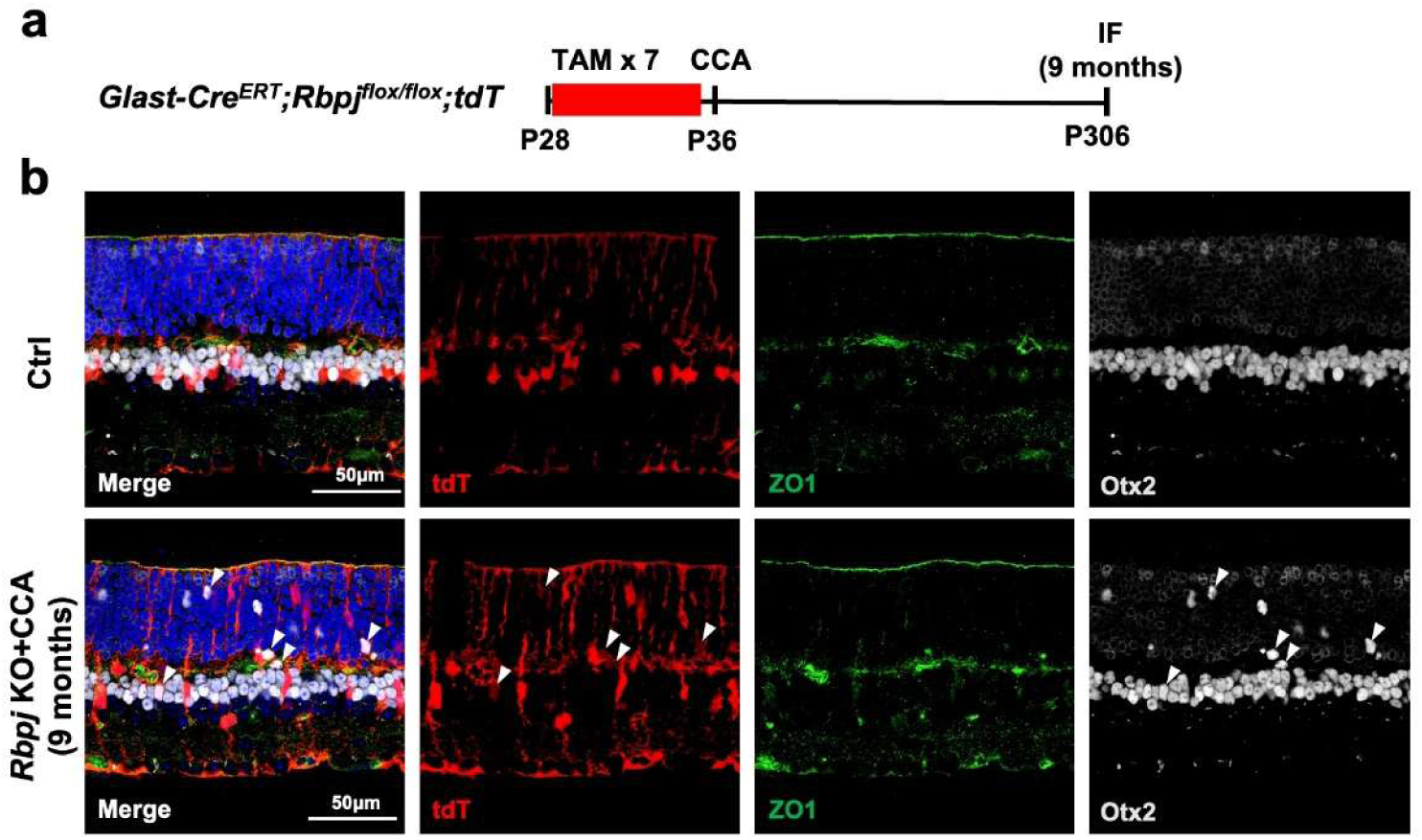
Expression pattern of ZO1 in the retina. (a) Schematic illustration of ZO1 staining experiment. (b) Representative immunostaining of Otx2 and ZO1 on retinal sections from *Glast-Cre^ERT^;tdT* and *Glast-Cre^ERT^;Rbpj^flox/flox^;tdT* mice at 9 months post TAM injection. The white arrows refer to tdT+ Otx2+ cells.

**Figure S23.**
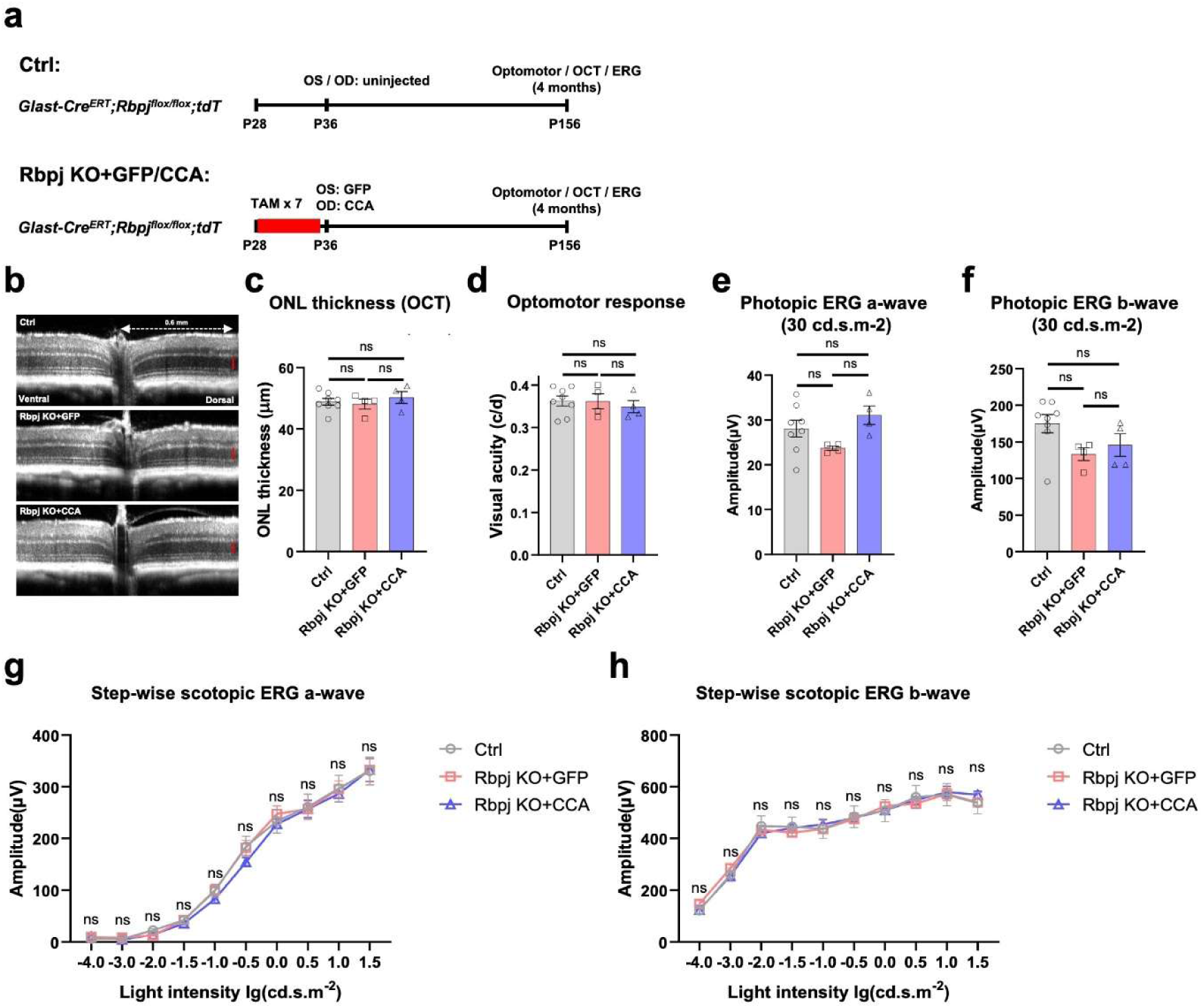
MG reprogramming induced by *Rbpj* deletion and CCA does not impair retinal structure and function. (a) Schematic illustration of experiment design for testing retinal structure and function. (b) Representative OCT images of retinas that were untreated or injected with GFP and CCA. (c) ONL thickness of retinas with different treatments. Each data point represents one mouse eye, data are presented as mean ± SEM. ns=not significant, by one-way ANOVA with Tukey’s post hoc test. (d) Optomotor test of mice with different treatments. (e-f) a- and b-wave photopic ERG response. light intensity 30 cd.s.m-2 Each data point represents one mouse eye, data are presented as mean ± SEM. ns=not significant, by one-way ANOVA with Tukey’s post hoc test. (g-h) a- and b-wave amplitudes of step-wise scotopic ERG responses of mice eyes by light intensity from -4.0 lg(cd.s.m-2) to 1.5 lg(cd.s.m-2). Each data point represents one mouse eye, data are presented as mean ± SEM. ns=not significant, by two-way ANOVA with Tukey’s post hoc test.

